# Beta-burst dynamics in the motor cortex are reshaped through sensorimotor refinement

**DOI:** 10.1101/2025.09.22.677717

**Authors:** Kate Schipper, Mahmoud Hassan, Bogdan Draganski, Paolo Ruggeri, Jérôme Barral

## Abstract

Beta-band activity (15–30 Hz) in motor cortex is closely linked to movement-related processing, yet its transient burst dynamics during long-term learning remain poorly understood. We used High-density electroencephalography (HD-EEG) to examine how beta bursts change over nine sessions of a bimanual coordination task, under adaptive and non-adaptive training conditions. Both training conditions led to motor skill learning, with the adaptive group improving more during training, but the non-adaptive group showing better retention. With training, contralateral primary motor cortex showed stronger beta desynchronization during movement and greater synchronization after movement. These changes reflected underlying burst dynamics: post-movement bursts became more temporally confined and consistent, with increased probability and reduced timing variability across sessions. Only the adaptive group showed a session-related increase in burst amplitude. These results demonstrate that beta burst features reorganize with practice, providing a temporally precise neural readout of training progression and revealing how different learning conditions shape cortical dynamics over time.

## Introduction

Motor skill learning relies on the coordinated activity of cortico-thalamo-basal ganglia circuits, which adjust with practice to enable more efficient and precise motor control (Hardwick et al., 2018; Park et al., 2022; Y. Peng & Wang, 2024). At the macroscopic level, these adjustments are partially reflected in modulations of neural oscillations—rhythmic patterns of activity measurable with non-invasive recordings such as EEG (Buzsáki & Draguhn, 2004; Hari & Salmelin, 1997; Jee, 2021). Oscillations emerge from the interplay of excitatory and inhibitory processes within these circuits, and long-term changes in this balance during learning can manifest as gradual shifts in oscillatory dynamics, making them a sensitive marker of training-related plasticity. Beta-band oscillations (15–30 Hz) are particularly prominent in motor regions and are closely linked to movement preparation, execution, and termination (Jasper & Penfield, 1949; Penfield, 1954; J. Peng et al., 2024). Before and during movement, beta power typically decreases—a phenomenon known as event-related desynchronization (ERD)—reflecting a release of cortical inhibition and increased motor readiness (Engel & Fries, 2010; Pfurtscheller, 2001; Pfurtscheller & Lopes Da Silva, 1999). ERD magnitude often increases with training, though such changes have been primarily documented within single practice sessions rather than across multiple training days (Moisello et al., 2015; Nelson et al., 2017; Ricci et al., 2019). ERD is also enhanced when upcoming actions are predictable, whether temporally or contextually (Alegre et al., 2003; Teodoro et al., 2018). After movement, beta power rebounds in a response known as event-related synchronization (ERS), which is thought to reflect the reinstatement of inhibitory control and ongoing performance monitoring (Cassim et al., 2001; Engel & Fries, 2010; Pfurtscheller, 1992; Spitzer & Haegens, 2017). ERS amplitude tends to increase as performance stabilizes and errors decline, potentially indexing confidence in internal models of action (Haar & Faisal, 2020; Porter et al., 2024; Qi & Qi, 2022; Tan et al., 2014, 2016; Torrecillos et al., 2015).

While ERD and ERS serve as reliable markers of motor system (dis)engagement (Espenhahn et al., 2017), most evidence of their modulation comes from studies using simple, well-practiced tasks that demand little learning, such as button presses or isolated self-paced contractions. In contrast, motor skill acquisition typically unfolds over time, requiring repeated attempts, error correction, and internal model updates (Arpin et al., 2017; Miall & Wolpert, 1996). Far less is known about how beta dynamics adapt during more complex forms of learning that require extended practice. Only a few studies have addressed this question directly. For instance, Jochumsen et al., (2017), observed an initial increase in ERD following one session of laparoscopic training, but no further change across six additional sessions. However, EEG was recorded during a separate probe task involving palmar grasps, which may not have fully engaged the trained motor circuits. More recently, Feng et al., (2024) tracked beta ERD over eight days of visuomotor force-tracking in three groups performing low, medium or high difficulty versions of the task. Difficulty was manipulated by varying the force control demands and the complexity of the target pattern. Across all groups, beta ERD in the dorsolateral prefrontal cortex (DLPFC) followed an inverted U-shaped trajectory, initially increasing, peaking and then returning toward baseline, a pattern that the authors interpret as in-line with the expansion–renormalization model of neuroplasticity (Lövdén et al., 2010; Wenger et al., 2017). Interestingly, this trajectory varied with task difficulty, with more demanding conditions prolonging the expansion phase before renormalization. Yet their task involved fixed difficulty and continuous visual feedback, potentially enabling early strategy stabilization and reducing demands for further adaptation.

Across these studies, ERD modulations appear strongest during the early stages of training and plateau thereafter. In fixed-difficulty designs, this pattern could lead to the interpretation that neural adjustments occur only early on. However, such design constraints may limit opportunities for continued exploration and refinement, potentially masking or leading to misinterpretation of longer-lasting modulations that might emerge if task demands were allowed to evolve with the learner’s skill level. As highlighted by the Challenge Point Framework (Guadagnoli & Lee, 2004) when task demands are not calibrated to the learner’s evolving skill level, errors and feedback lose informational value. Practice becomes either too easy to drive adaptation or too punishing to support it, reducing the likelihood that sustained adjustments will emerge. This possibility motivated our comparison of adaptive and non-adaptive training, to test whether scaling task difficulty with performance modulations could sustain exploration and drive continued neural adjustment across sessions.

Recent evidence suggests that conventional time–frequency averaging may, in fact, overlook important features of motor-related beta activity that underlie these ERD and ERS patterns (Wessel, 2020). Rather than sustained oscillations, beta activity appears to comprise brief, high-amplitude bursts whose timing and probability vary across trials (Feingold et al., 2015; Lundqvist et al., 2024; Sherman et al., 2016; van Ede et al., 2018). These bursts, shaped by laminar-specific inputs within motor cortex, may underpin the trial-specific flexibility needed for skilled behaviour (Jones, 2016; Seedat et al., 2020a). From this perspective, ERD and ERS likely reflect shifts in burst probability rather than sustained amplitude changes. Notably, the timing of beta bursts—particularly in pre- and post-movement periods—is more tightly associated with behavioural outcomes than averaged beta power, with early bursts linked to faster responses and later bursts associated with error detection and post-performance adjustment (Little et al., 2019; Wessel, 2020). The demands on this trial-by-trial neural flexibility are likely shaped by the learning conditions. Over extended practice, both adaptive and non-adaptive learners may show reorganization of burst dynamics—such as changes in burst probability during key task periods and reduced timing variability across trials—as they acquire the skill. However, in adaptive training, where task demands evolve and motor output is continually refined, these adjustments may be more pronounced, producing sharper and more precise fine-tuning of when and how often bursts occur. Tracking how burst dynamics change with practice could thus yield a more fine-grained view of neural adaptation, complementing traditional metrics and revealing how the motor system becomes more structured and efficient over time (Shin et al., 2017; Torrecillos et al., 2018). However, just as most investigations of ERD and ERS have been limited to simple tasks or short-term training, research on beta bursts has also focused almost exclusively on single-session paradigms with minimal learning demands. As a result, it remains unknown how beta bursts reorganize with extended practice, or how their dynamics relate to traditional ERD/ERS patterns when motor behaviour must be continuously refined over time.

To directly address this gap, the present study examines both trial-averaged (ERD/ERS) and burst-resolved beta dynamics across multiple sessions of a complex bimanual force coordination task, which required participants to maintain a steady force with one hand while producing precisely timed ballistic output with the other. This combination of sustained and time-critical control made the task inherently difficult, requiring participants to explore and refine their motor performance across trials, engaging flexible sensorimotor control rather than relying on fixed response strategies. In the adaptive group, task difficulty increased as performance improved, sustaining the need for ongoing motor exploration and refinement across sessions. The non-adaptive group performed the same task at constant difficulty. EEG was recorded during early (Session 1), middle (Session 5), and late (Session 9) phases of training (see Fig. 1), enabling us to track how neural dynamics evolved over time.

**Figure 1.**
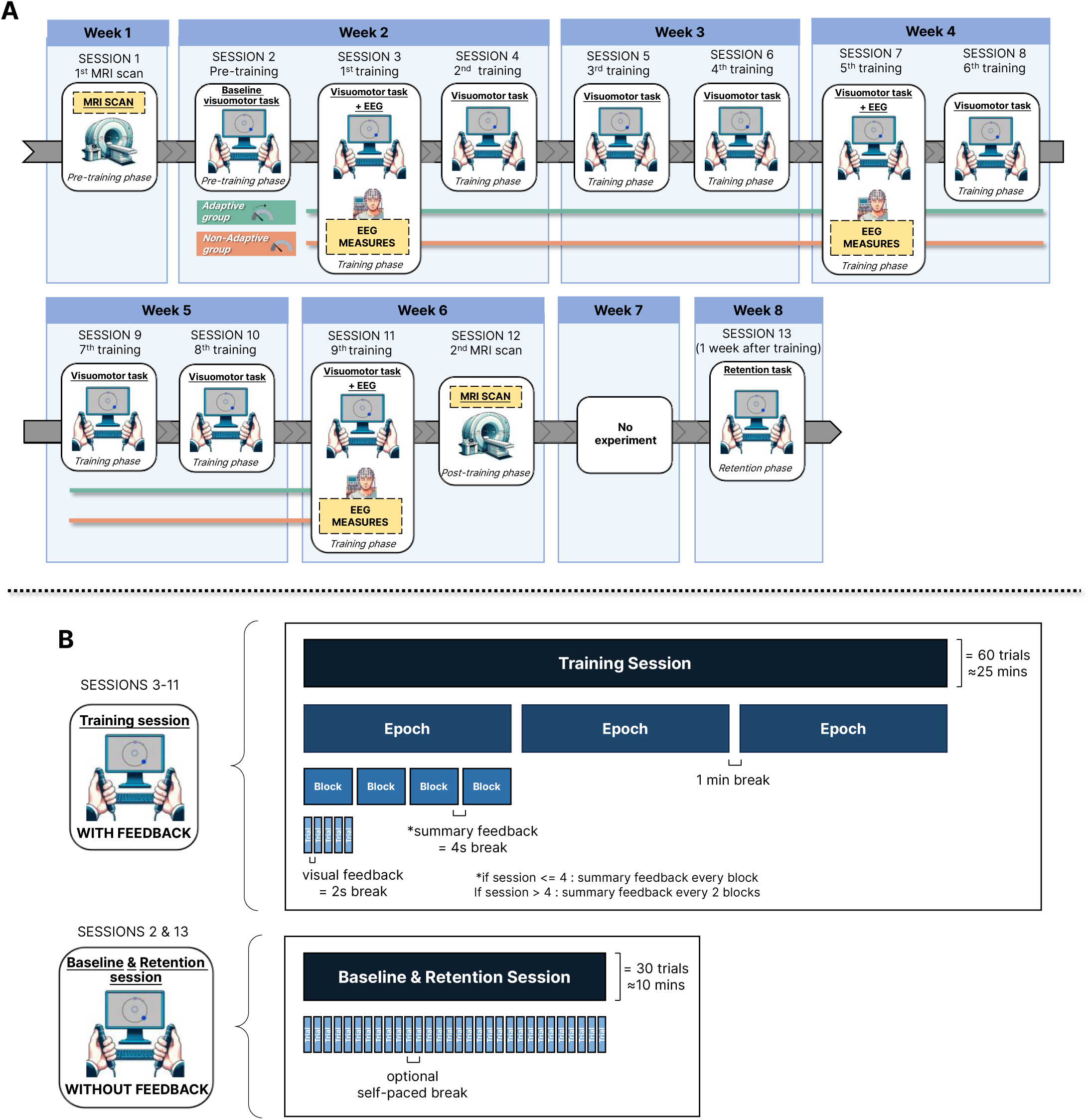
Protocol and Experimental design. The protocol spanned 13 sessions across 8 weeks, including MRI (Sessions 1 and 12) and EEG recordings (Sessions 3, 7, and 11). Participants completed visuomotor training from Sessions 4 to 12. Each training session consisted of 3 epochs of 4 blocks, with 5 trials per block (60 trials total). Including breaks, each session lasted approximately 25 minutes. Baseline (Session 2) and retention (Session 13) sessions were conducted without feedback. Participants were assigned to either an adaptive or non-adaptive training group.

We began by analyzing beta-band activity at the sensor level to identify regions showing learning-related changes. Source localization then guided our focus to bilateral primary motor cortex (M1), where beta dynamics were most prominent. At the behavioural level, we expected (1) general improvements in motor performance across sessions, as participants refined coordination of the bimanual task; and (2) greater and more sustained improvements in the Adaptive group compared to the Non-Adaptive group, due to the incremental demands placed on motor control when difficulty scaled with performance. At the neural level, our hypotheses were threefold: (1) we expected ERD and ERS amplitudes to increase with practice, particularly in right M1. While previous studies often report that such changes peak early or remain minimal across extended training, the sustained demands of our complex task—and especially the adaptive condition, where difficulty scaled with performance—were expected to prolong cortical adjustment and yield measurable changes across sessions. (2) We hypothesized that these average power changes would reflect greater temporal segregation of beta bursts across task phases. Specifically, bursts were expected to occur less frequently during movement (reflecting disinhibition and readiness) and more frequently after movement (reflecting inhibitory resetting and performance monitoring). (3)

Finally, we anticipated that burst probability, timing, and amplitude would reorganize more strongly under adaptive training conditions, due to the sustained need for motor refinement. By combining high-resolution EEG, burst-resolved metrics, and a longitudinal design, this study aimed to clarify how beta activity reorganizes with practice to support motor skill learning under different training conditions.

## Results

We began by examining the temporal profile of motor output across the three EEG sessions to ensure that movement execution was comparable between sessions. We then assessed behavioural performance from baseline to retention to evaluate learning. Subsequent analyses focused on neural activity, examining beta-band dynamics at the scalp and in source-localized motor cortex, and characterising transient beta bursts in terms of their probability, timing variability, and amplitude. All neural analyses targeted the bimanual moment of the task, using a time window centered on the peak force of the left hand.

### Motor output becomes more consistent with practice

We recorded grip force data from 31 participants (16 Adaptive, 15 Non-Adaptive) as they completed a bimanual coordination task. The protocol began with a baseline session of 30 no-feedback trials, followed by nine training sessions of 60 trials each, and ended with a retention session of 30 no-feedback trials. In each trial, participants maintained steady right-hand force while producing a brief, precisely timed left-hand pulse to shoot a visual target. This left-hand action— performed while the right hand sustained constant force—represents the most demanding phase of sensorimotor coordination. We therefore centered our analysis of motor output and neural activity on the time window surrounding the left-hand shot (Fig. 2B, panel 4).

**Figure 2.**
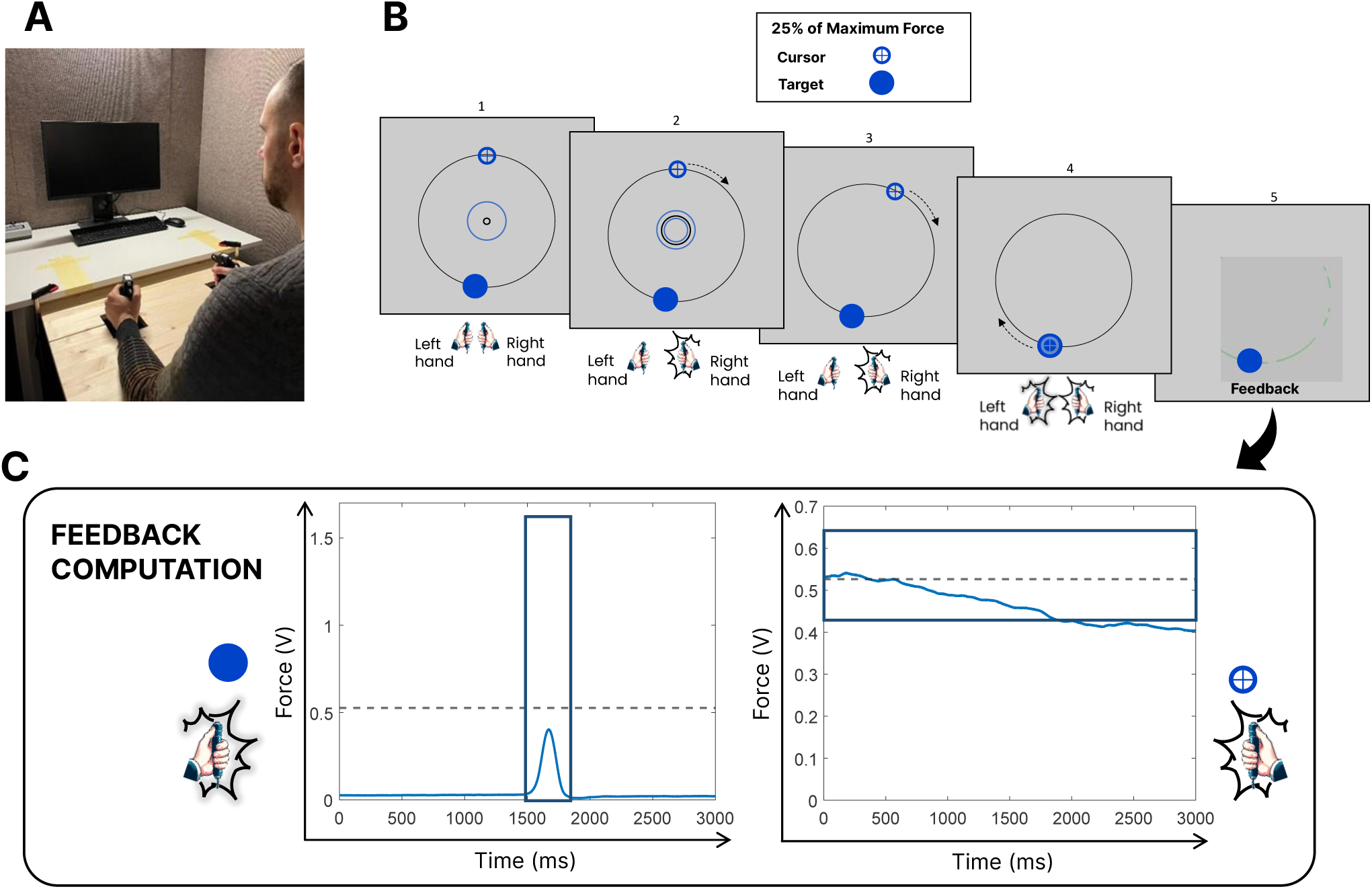
Bimanual force-coordination task. (A) Experimental set-up. Participants performed the task using two hand-held dynamometers – picture of one of the co-authors. (B) Experimental task: each trial began with a steady right-hand force (25% MVC) held within a threshold zone for 300 ms (B, panel 2). Once stable, the feedback disappeared, and a cursor completed a 3-second circular trajectory. When the cursor reached the target, participants applied a brief left-hand force pulse (25% MVC) to shoot. Feedback was provided after each trial based on whether force remained within the tolerance ranges (blue boxes in panel C). (C) Example force traces: left-hand (left) and right-hand (right) force profiles from a single trial.

Visual inspection of left-hand force traces revealed systematic changes in the shape of motor output with practice across both groups (Fig. 3). In Session 1, adaptive participants tended to produce more sharply peaked, time-locked responses, while non-adaptive participants showed greater variability, including slower ramps and prolonged force application. By Session 5, both groups exhibited more refined, phasic force profiles, with the adaptive group maintaining slightly steeper onsets. In Session 9, this overall pattern persisted, though some adaptive participants displayed increased variability, possibly reflecting adjustments to higher task difficulty introduced for some individuals at this stage.

**Figure 3.**
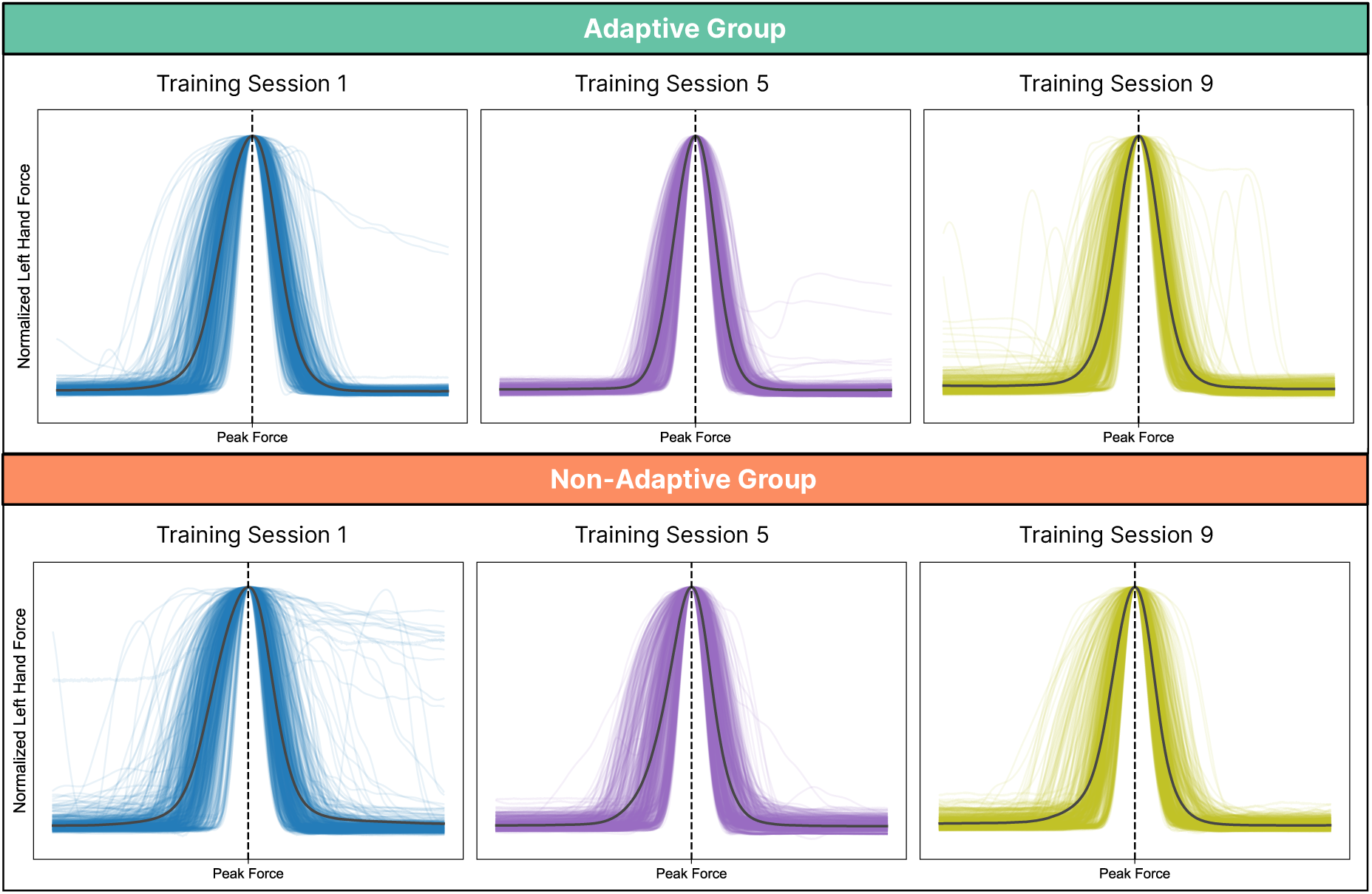
Normalized left-hand force profiles across training sessions. Force traces for Sessions 1 (blue), 5 (purple), and 9 (green) are shown for the adaptive group (top row) and non-adaptive group (bottom row). Thin lines represent individual trials from all participants; thick black lines indicate the average across trials. Force was normalized within each trial to a peak of 1 and time-aligned to peak force (dashed vertical line).

These qualitative trends were supported by statistical analysis of the area under the curve (AUC) for each trial. A generalized linear mixed model (GLMM) revealed a significant main effect of Session (χ^2^(2) = 695.35, *p* < .001) and a significant Session × Group interaction (χ^2^(2) = 30.94, *p* < .001), indicating that the trajectory of AUC changes differed between groups. The main effect of Group was not significant (χ^2^(1) = 0.86, *p* = .353). AUC significantly decreased from Session 1 to Sessions 5 and 9 in both groups (*all p* < .001), reflecting increasingly phasic, temporally focused motor responses. Importantly, the adaptive group showed a significant increase in AUC from Session 5 to Session 9 (*z* = −7.92, *p* < .001), consistent with the observed increase in variability and likely related to the increased task difficulty for some individuals in this session.

To account for session-related changes in left-hand response profiles, neural analyses were time-locked to the peak force, the culmination of the motor action and the most behaviourally consistent reference point. This alignment helped minimize variability across trials related to differences in movement onset, duration, and shape.

Although our analyses focus on the time window surrounding the left-hand shot, right-hand force traces across the full 3-second trial are shown in Supplementary Fig. S1 to illustrate how force maintenance performance evolved throughout training.

### Adaptive Training Leads to Greater Improvements in Bimanual Motor Performance but Performance Gains Are Not Fully Retained

To quantify overall performance in the bimanual task, we used a composite metric called the Bimanual Performance Deviation Score (BPDS). This score captured trial-wise deviations across three dimensions: (1) right-hand force stability during the 226 ms period preceding the left-hand peak force, (2) left-hand peak force, and (3) timing error, defined as the temporal deviation between the peak of the left-hand force and the center of the visual target. Each component was expressed as a deviation from its target value, and the three were combined into a single Euclidean distance metric. The resulting BPDS provided a continuous measure of coordination accuracy, with lower scores indicating better performance.

To assess motor learning and the retention of these gains, we ran a single GLMM including all three key sessions (Baseline, last training - Session 9, and Retention), with Group (Adaptive, Non-Adaptive) and Session as fixed factors and Participant as a random factor. The model revealed a significant main effect of Session (χ^2^(2) = 2504.60, p < .001), a non-significant main effect of Group (χ^2^(1) = 0.01, p = .916), and a significant Group × Session interaction (χ^2^(2) = 10.21, p = .006).

For the Baseline–Retention comparison, post hoc tests (Bonferroni-corrected) showed significant reductions in BPDS for both groups (Adaptive: z = 31.86, p < .001; Non-Adaptive: z = 30.99, p < .001). In the Adaptive group, BPDS decreased from 2.70 ± 0.93 at PRE to 1.55 ± 0.90 at Retention. The Non-Adaptive group showed a similar pattern, with BPDS declining from 2.44 ± 0.77 to 1.62 ± 1.05. No significant differences between groups at baseline (z = 0.40, p = 1.000) or at retention (z = −0.22, p = 1.000), confirm that the interaction effect reflects a difference in the magnitude of change rather than a difference in final performance.

For the Last training-Retention comparison, post hoc tests (Bonferroni-corrected) revealed a small but significant increase in BPDS for the Adaptive group (z = −4.85, p < .001), whereas no significant change was observed for the Non-Adaptive group (z = −0.29, p = 1.000). BPDS in the Adaptive group rose from 1.41 ± 0.85 at Session 9 to 1.55 ± 0.90 at Retention, while in the Non-Adaptive group it increased only slightly from 1.58 ± 1.06 to 1.62 ± 1.05, indicating that performance was maintained (see Fig. 4).

**Figure 4.**
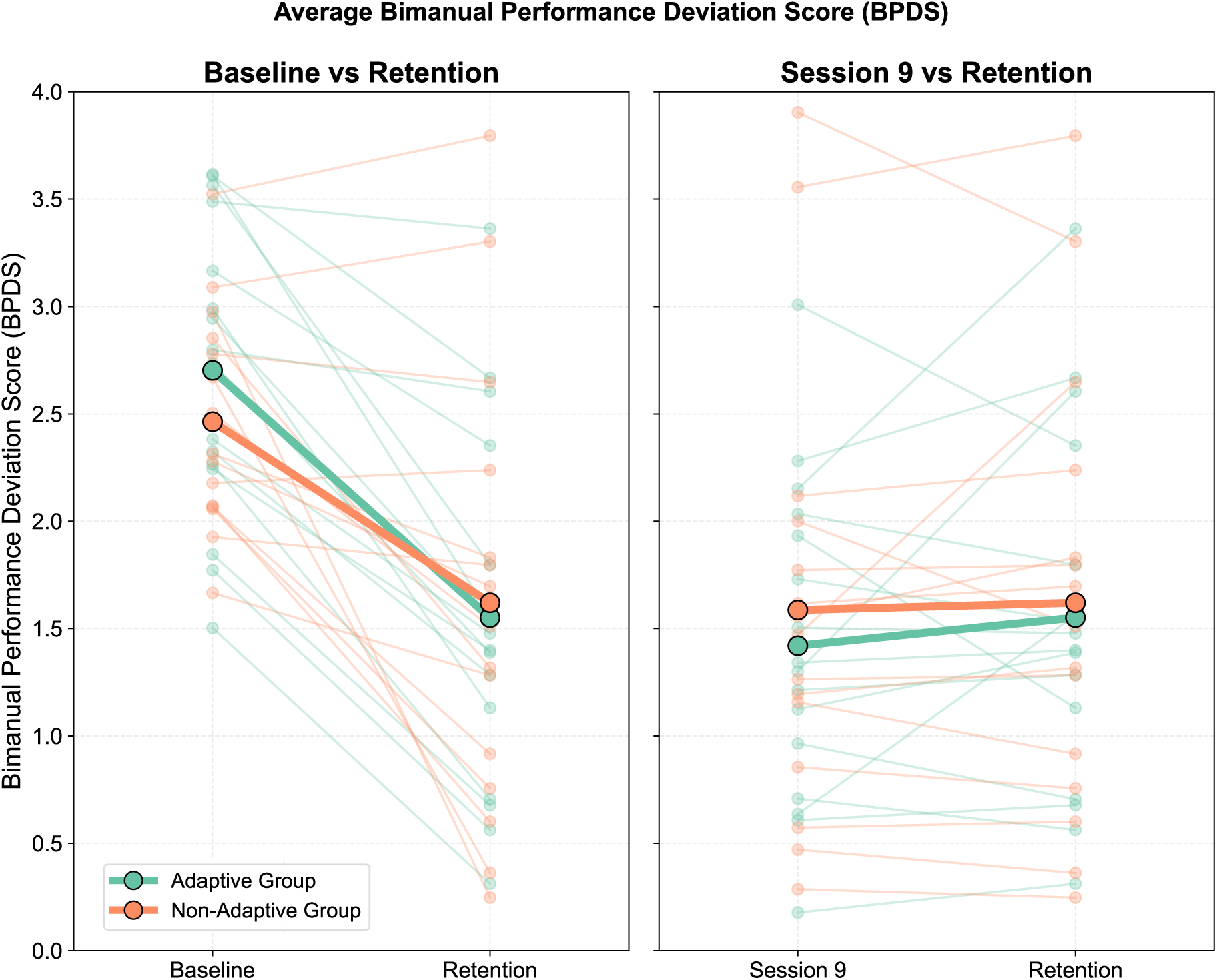
Bimanual Performance Deviation Scores (BPDS) for individual participants in the Adaptive (teal) and Non-Adaptive (orange) groups. Lower values indicate more accurate bimanual coordination. Left panel: Scores from Baseline and Retention sessions. Right panel: Scores from Session 9 and Retention. Each line represents a single participant; bold lines indicate group means.

### Scalp-Level Beta Dynamics : Session-Related Changes in Oscillatory Power

#### Post-Movement Beta Synchronization Increases With Training

To contextualize our targeted beta-band analysis, we first examined broadband time– frequency activity (4–40 Hz) to visualize the full spectral profile of movement-related responses. While some modulation was present at lower frequencies (e.g., lower alpha range), activity was most prominent and topographically consistent in the beta band (15-30 Hz). Full spectral maps of Session 1 and 3 are provided in Supplementary Fig. S2.

We then analyzed beta-band activity to assess training-related changes in neural dynamics. Importantly, no significant group × session interaction effects were observed at either the scalp or source level, so data were pooled across groups. At the scalp level, sensor-level time–frequency analysis revealed a significant increase in post-movement beta synchronisation between EEG Session 1 and Session 3. This appeared as a single negative cluster in the statistical contrast (p = 0.044), over contralateral centro-parietal electrodes around 1000 ms after peak force, with the strongest effects at electrodes B21, B22, and B23 (located approximately over C2, C4, and C6 in the 10–20 system). Time courses at these sites confirmed enhanced beta power during the post-movement period in EEG Session 3 (Fig. 5, bottom). A brief beta peak at movement onset in Session 1 likely reflects muscle-related artifact in four participants, as suggested by concurrent high-frequency activity in the broadband spectrograms (Supplementary Fig. S2). For full topographies and session-wise time–frequency maps in beta band, see Supplementary Fig. S3.

**Figure 5.**
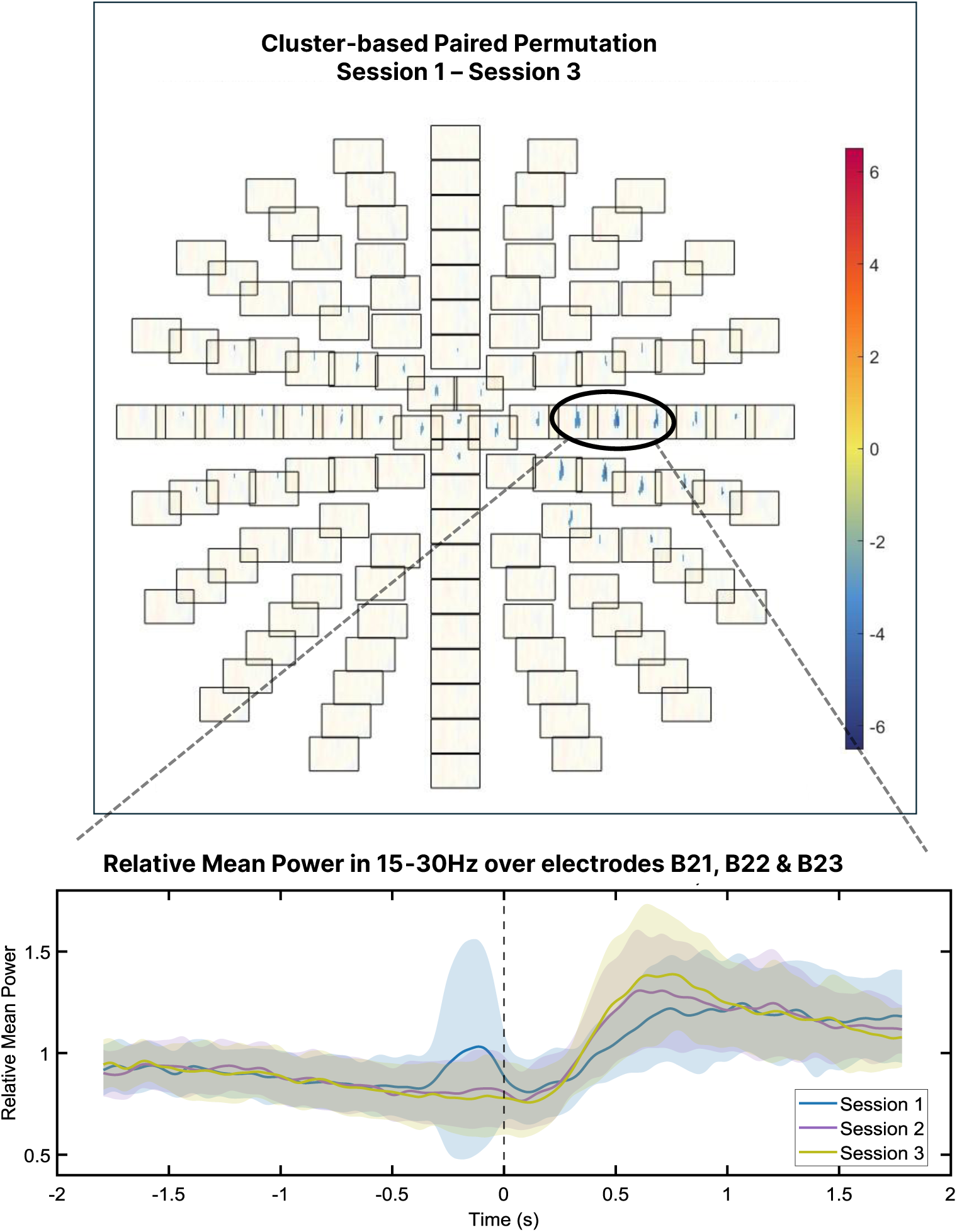
Sensor-level differences in beta-band activity between Session 1 and Session 3. Top: Topographical t-map from a cluster-based permutation test comparing beta-band (15–30 Hz) power between Session 1 and Session 3 across the −0.75 to +1 s window around movement onset. Each box represents an electrode, and colours indicate t-values. Blue tones reflect lower beta power in Session 1 relative to Session 3. Bottom: Time courses of mean relative beta power (±1 SD) averaged across the significant electrodes (B21, B22, B23), aligned to movement onset (0 s, dashed line). A larger post-movement beta rebound is evident in Session 3. A brief beta peak around movement onset in Session 1 reflects movement-related artifact in a subset of participants.

### Source-Level Beta Dynamics : Cortical Localization of ERD/ERS Changes

#### Training Enhances Beta Desynchronization and Rebound in Right Sensorimotor Cortex

To better localize the scalp-level effects and identify their cortical generators, we analyzed source-level beta power in EEG Sessions 1 and 3.

Cluster-based paired permutation tests were performed on source-level beta power across two task-relevant windows: the movement period (−400 to +250 ms) and the post-movement period (+250 to +1000 ms). During the movement window, a significant positive cluster (p = 0.006) was observed over right sensorimotor cortex, indicating stronger beta suppression in Session 3 compared to Session 1 (red tones indicate higher beta power in Session 1; blue tones indicate higher beta power in Session 3; see Figure 6). During the post-movement period, a significant negative cluster (p = 0.020) over the same region indicated enhanced beta rebound with training. No significant group × session interaction was found in either window, suggesting that training-related changes in cortical beta dynamics occurred similarly in both groups (see Supplementary Fig. S4 for full time-resolved maps).

**Figure 6.**
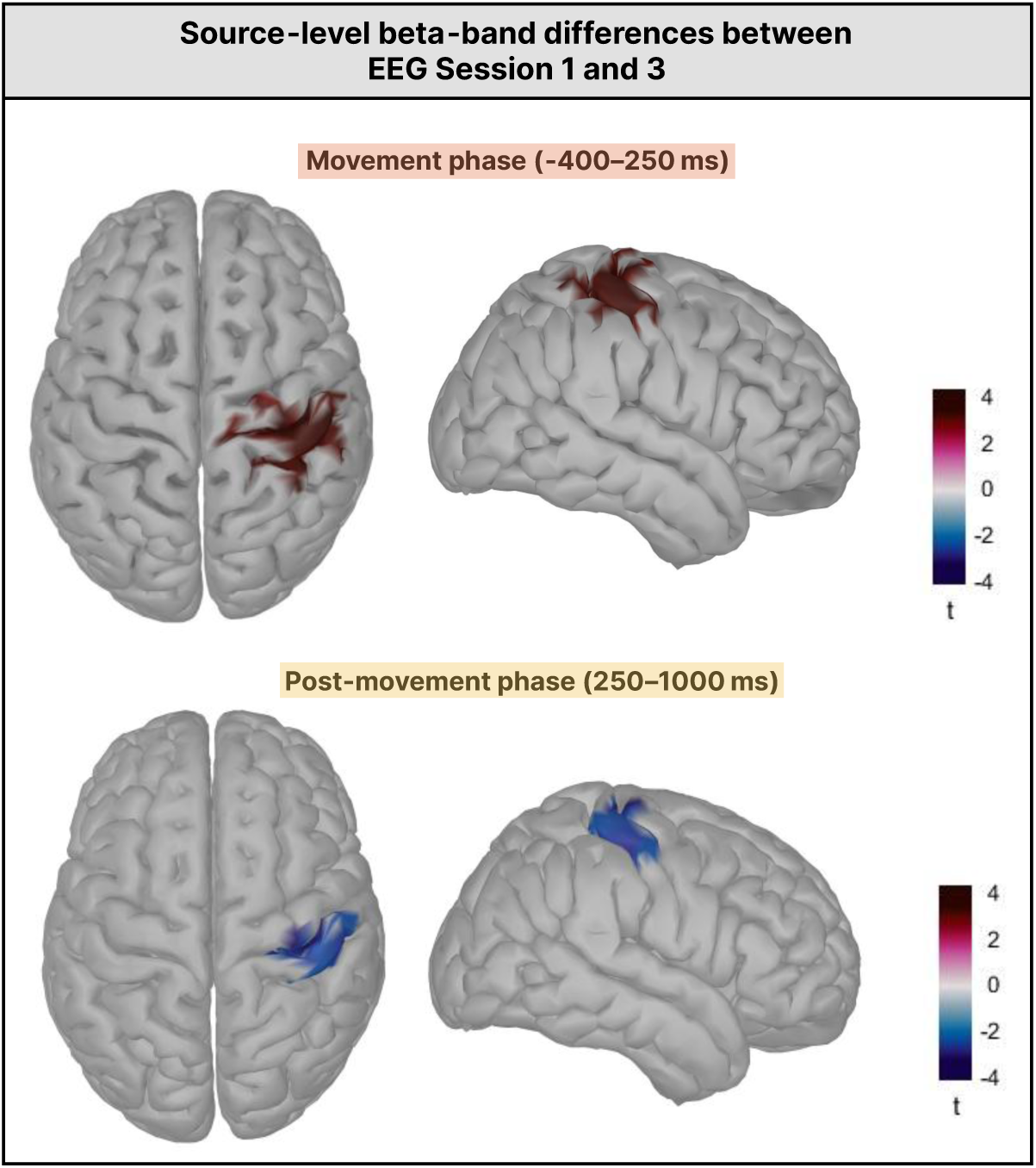
Source-level beta-band power differences between Session 1 and Session 3 during movement and post-movement periods. Top row: *t*-maps showing the statistical contrast of beta power between sessions during the movement window (−400 to +250 ms relative to movement onset). Bottom row: *t*-maps for the post-movement window (250 to 1000 ms). Cortical surfaces are shown in dorsal and right lateral views. Red tones indicate higher beta power in Session 1; blue tones indicate higher beta power in Session 3.

As observed at the scalp level, a brief beta peak at movement onset in Session 1 likely reflected muscle-related artefacts. Visual inspection of the broadband spectrograms confirmed that these peaks coincided with high-frequency activity (See Fig. S2). At the source level, these artefacts appeared as lateralised patches of high beta power, distinct from the bilateral sensorimotor clusters identified in the statistical analysis (Fig. S4), indicating that the reported training-related changes were not driven by artefacts.

### Beta Burst Dynamics : Session-Related Changes in Probability, Amplitude and Timing

The session-related changes in beta power described above likely reflect alterations in the occurrence of transient beta bursts—brief, high-amplitude events that give rise to classical ERD and ERS patterns. We next examined how these bursts in M1 evolved with training, focusing on their temporal distribution, probability of occurrence, and amplitude. Beta bursts were detected using an empirically optimized thresholding procedure, adapted from Little et al. (2019). The optimal threshold, defined as a number of standard deviations (SD) above the median beta amplitude, was determined by selecting the value that maximized the correlation between burst rate and mean amplitude across all subjects, sessions, and M1 scouts (see Materials and Methods). This procedure identified 1.4 SD above the median as the most effective threshold for burst detection in our data.

To visualize the timing of beta bursts across training, raster plots were generated for left and right M1, aligned to the left-hand shot (Fig. 7). This focus follows from the source-level results, which highlighted M1 as the main site of training-related beta power changes. In right M1, contralateral to the shooting hand, bursts became increasingly concentrated in the post-movement period over the course of training, with this pattern clearly emerging by Session 3. A transient suppression of bursts around movement onset also became more apparent across sessions. In contrast, burst timing in left M1 (ipsilateral to the shot, and associated with the stabilizing hand) was more diffusely distributed, though a modest post-movement increase emerged by Sessions 2 and 3.

**Figure 7.**
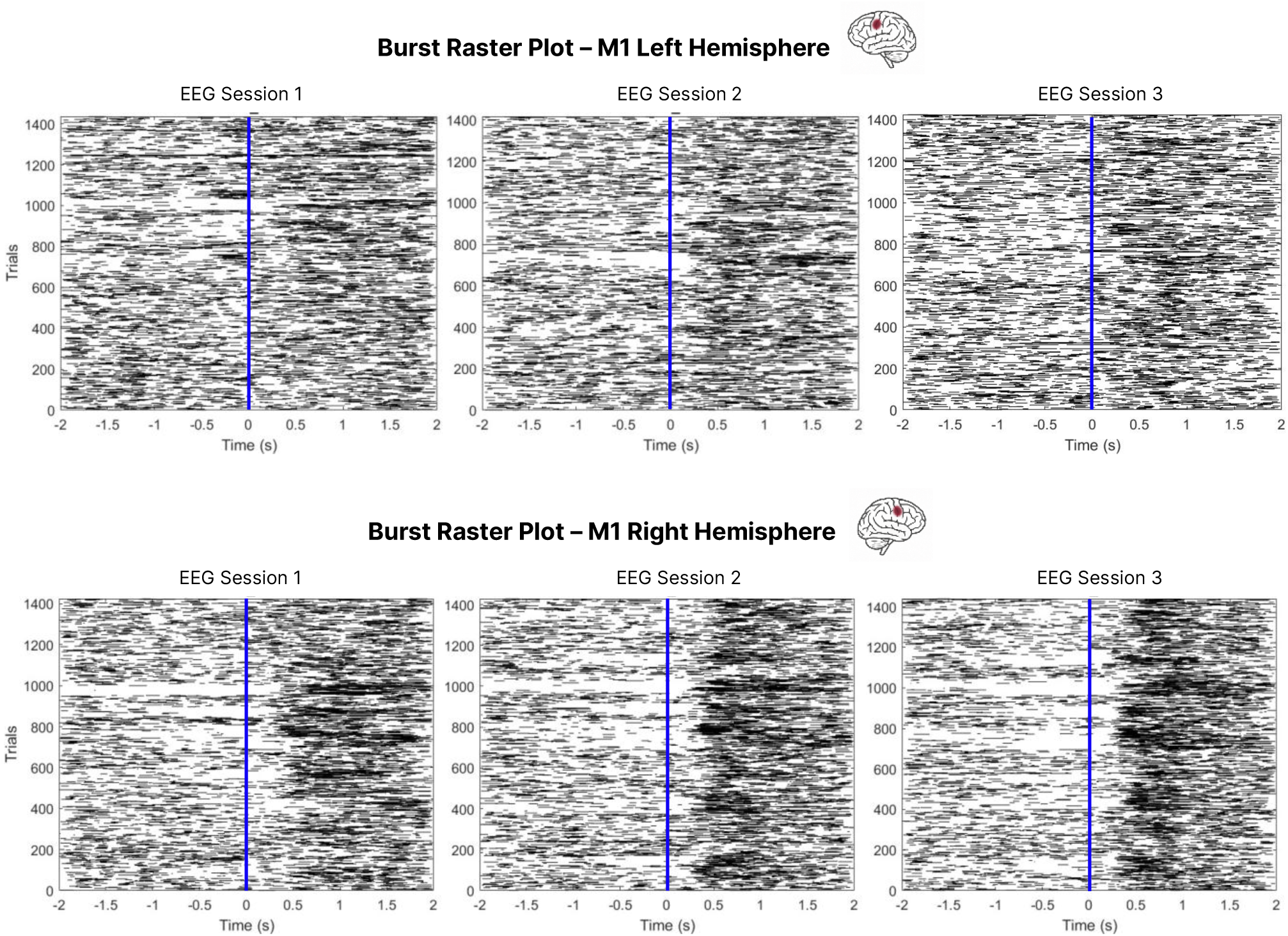
Temporal distribution of beta bursts across sessions and hemispheres. Raster plots show the timing of detected beta bursts relative to movement onset (vertical blue line at 0 s), across all trials and participants. Each row represents one trial; each black tick marks the time of a beta burst. Plots are organized by session (left to right: Sessions 1, 2, and 3) and hemisphere (top row: left M1; bottom row: right M1).

#### Training modulates beta burst probability during and after movement

To complement the raster plots, we analyzed time-resolved beta burst probability, defined as the proportion of trials in which a burst occurred at each time point (Fig. 8). This measure captures how the likelihood of bursting varies across the trial. To examine differences across task phases, we computed burst probability within three predefined time windows: Baseline (−1750 to −750 ms), Movement (−750 to 250 ms), and Post-Movement (250 to 1250 ms). Repeated-measures ANOVAs were conducted separately for left and right M1, with Time Window and Session (S1–S3) as within-subject factors and Group (Adaptive, Non-Adaptive) as a between-subject factor.

**Figure 8.**
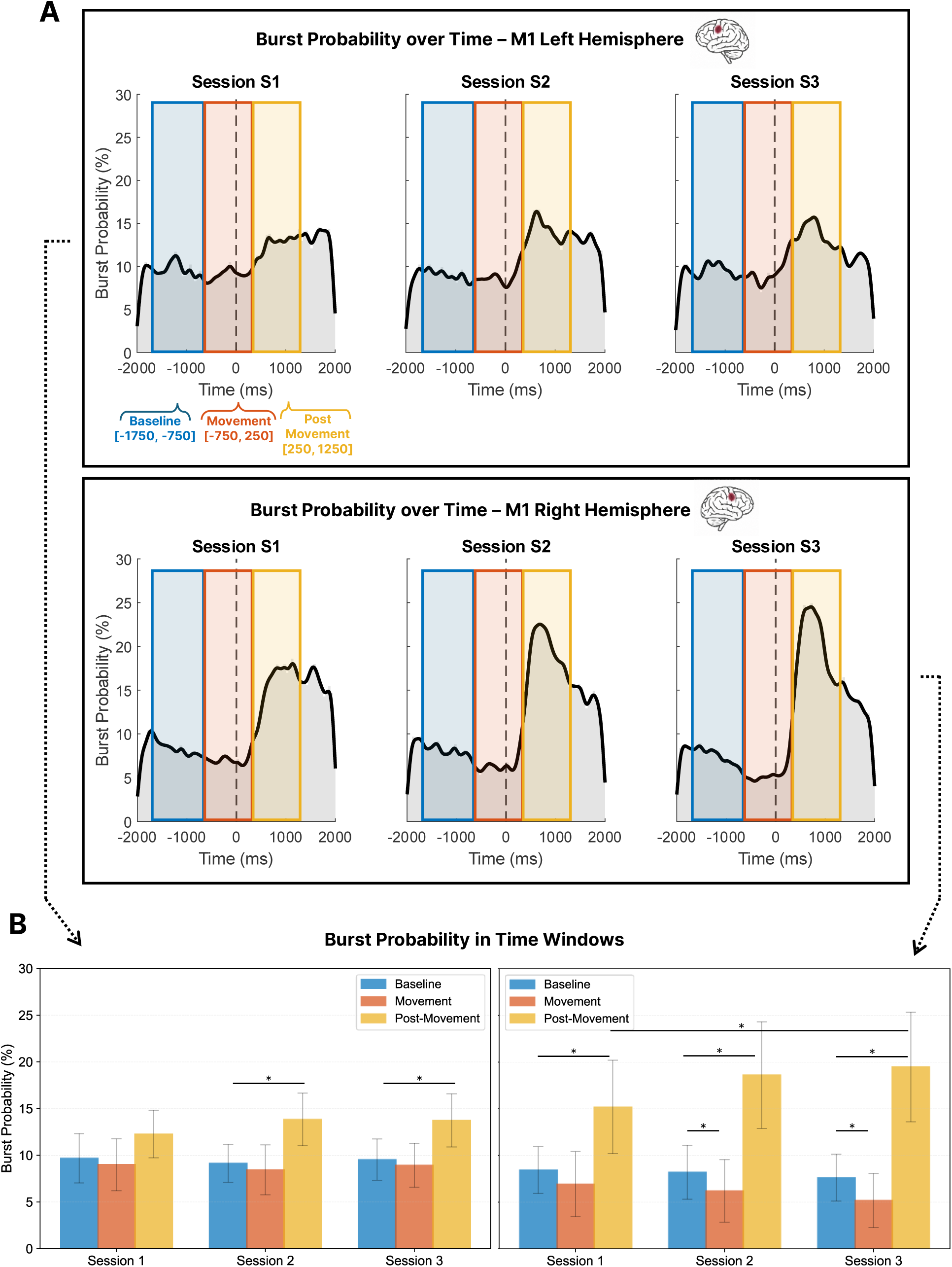
Beta burst probability across predefined time windows, sessions, and hemispheres. Time-resolved beta burst probability in right M1, averaged across all participants and shown separately for Sessions 1–3. Coloured rectangles indicate the three predefined analysis windows: Baseline (blue, −1750 to −750 ms), Movement (red, −750 to 250 ms), and Post-Movement (yellow, 250 to 1250 ms), aligned to movement onset (dashed vertical line at 0 ms). (B) Average beta burst probability (%) within each time window and session, plotted separately for left M1 (left panel) and right M1 (right panel). Asterisks indicate statistically significant differences (p < .05).

In right M1, we observed a robust main effect of Time Window (F(1.06, 26.53) = 51.19, *p*<.001, Greenhouse–Geisser corrected), indicating that burst probability differed across task phases. Post hoc Bonferroni-corrected comparisons showed that Post-Movement burst probability (mean across groups and sessions ≈ 17–20%) was significantly higher than both Baseline (≈ 7–9%) and Movement (≈ 5–7%). A significant Time Window × Session interaction was also found (F(1.57, 39.27) = 10.94, *p* < .001), reflecting training-related changes in the temporal profile of bursting. Post hoc tests (Bonferroni-corrected) showed that in all sessions, Post-Movement burst probability was significantly higher than both Baseline and Movement windows (*ps* < .001), and Baseline exceeded Movement in Sessions 2 and 3 (*ps* ≤ .004). Mean Post-Movement burst probability increased across sessions, from 15.1 ± 5.39% in Session 1 to 18.6 ± 5.97% in Session 2 and 19.4 ± 6.12% in Session 3; this increase was statistically significant from Session 1 to Session 3 (*p* = .009), consistent with an enhanced beta rebound over time. No main or interaction effects involving Group were observed (*ps* > .10). To check that the observed increase in post-movement burst probability was not driven by shifts in burst timing or window boundaries, we ran a complementary analysis using a single wide post force peak time window (0–2000 ms). This analysis confirmed a significant increase in burst probability from Session 1 to Session 3 (*p* = .031), indicating that the effect reflects a general amplification of post-movement bursting rather than a temporal shift.

In left M1, a similar analysis revealed a significant effect of Time Window (F(1.26, 31.47) = 24.18, *p* < .001). Post hoc comparisons confirmed that burst probability was significantly higher in the Post-Movement window compared to both Baseline (p < .001, Bonferroni-corrected) and Movement (p < .001, Bonferroni-corrected). The Time Window × Session interaction was also significant (F(4, 100) = 2.53, *p* = .045), indicating that the temporal structure of bursting evolved with training. Post hoc comparisons revealed that in Sessions 2 and 3, Post-Movement burst probability was significantly higher than both Baseline and Movement (*ps* < .01), though no significant differences were found between Baseline and Movement (p = .196). The main effect of Session was not significant (F(2, 50) = 1.82, *p* = .173) and no Group-related effects emerged.

#### Training Improves the Temporal Precision of Post-Movement Beta Burst Probability

To assess the temporal precision of post-movement beta activity, we quantified how consistently the peak in burst probability occurred across trials. For each participant and session, we calculated the standard deviation of the time points at which burst probability reached its maximum within the post-movement window. The average variability in peak burst probability timing decreased across sessions, from 467 ± 78 ms in Session 1, to 425± 81.5 ms in Session 2, and 410 ± 91.7 ms in Session 3 (see Fig.9).

**Figure 9.**
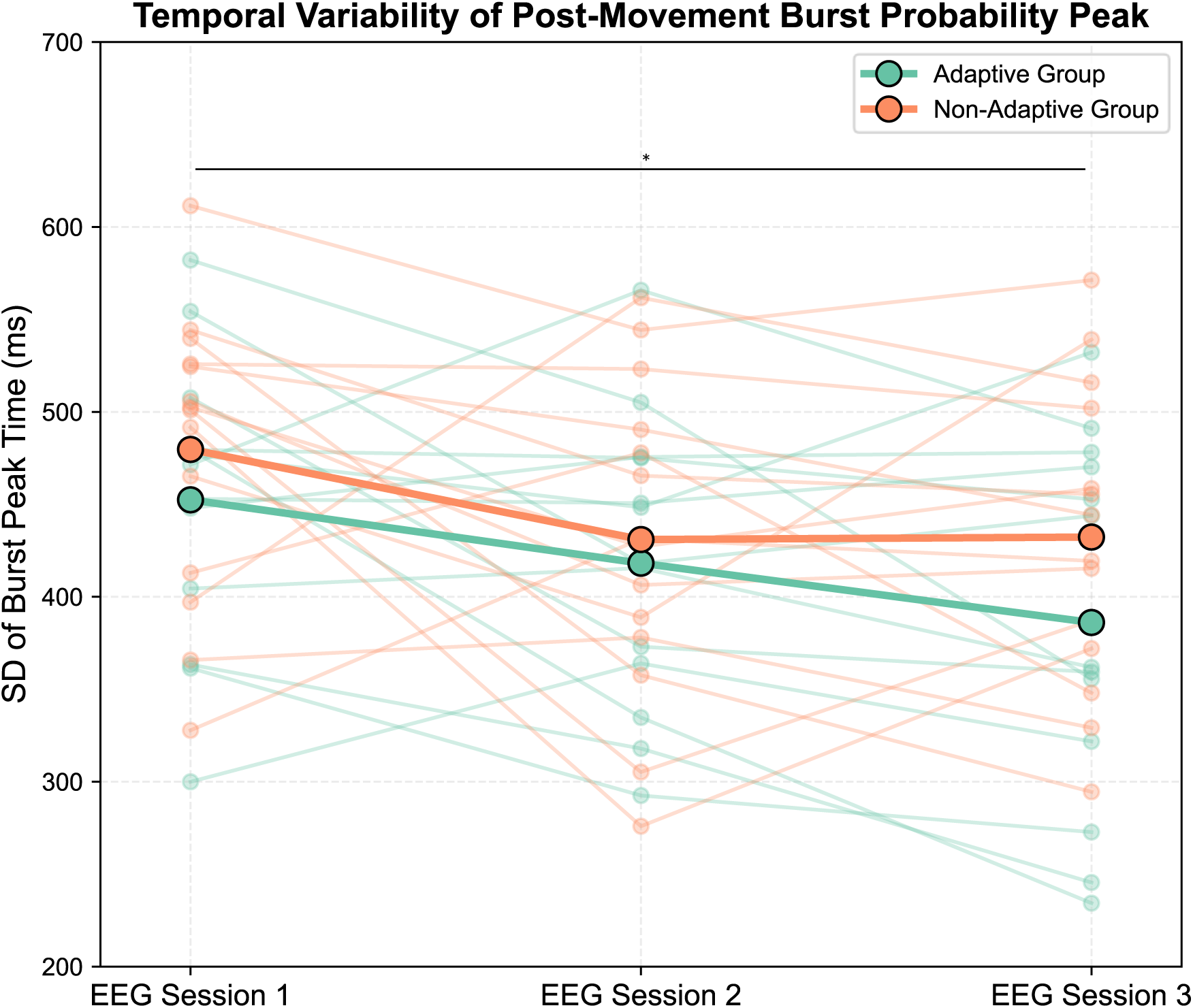
Temporal variability of post-movement burst peak probability across training sessions. Standard deviation of burst peak probability time (in milliseconds) is shown for each participant (faint lines and dots) in the Adaptive (teal) and Non-Adaptive (orange) groups across Sessions 1, 2, and 3. Bold lines and larger markers represent group means. A decrease in variability over time is visible in both groups Asterisks indicate timepoints where pairwise differences reached significance.

**Figure 10.**
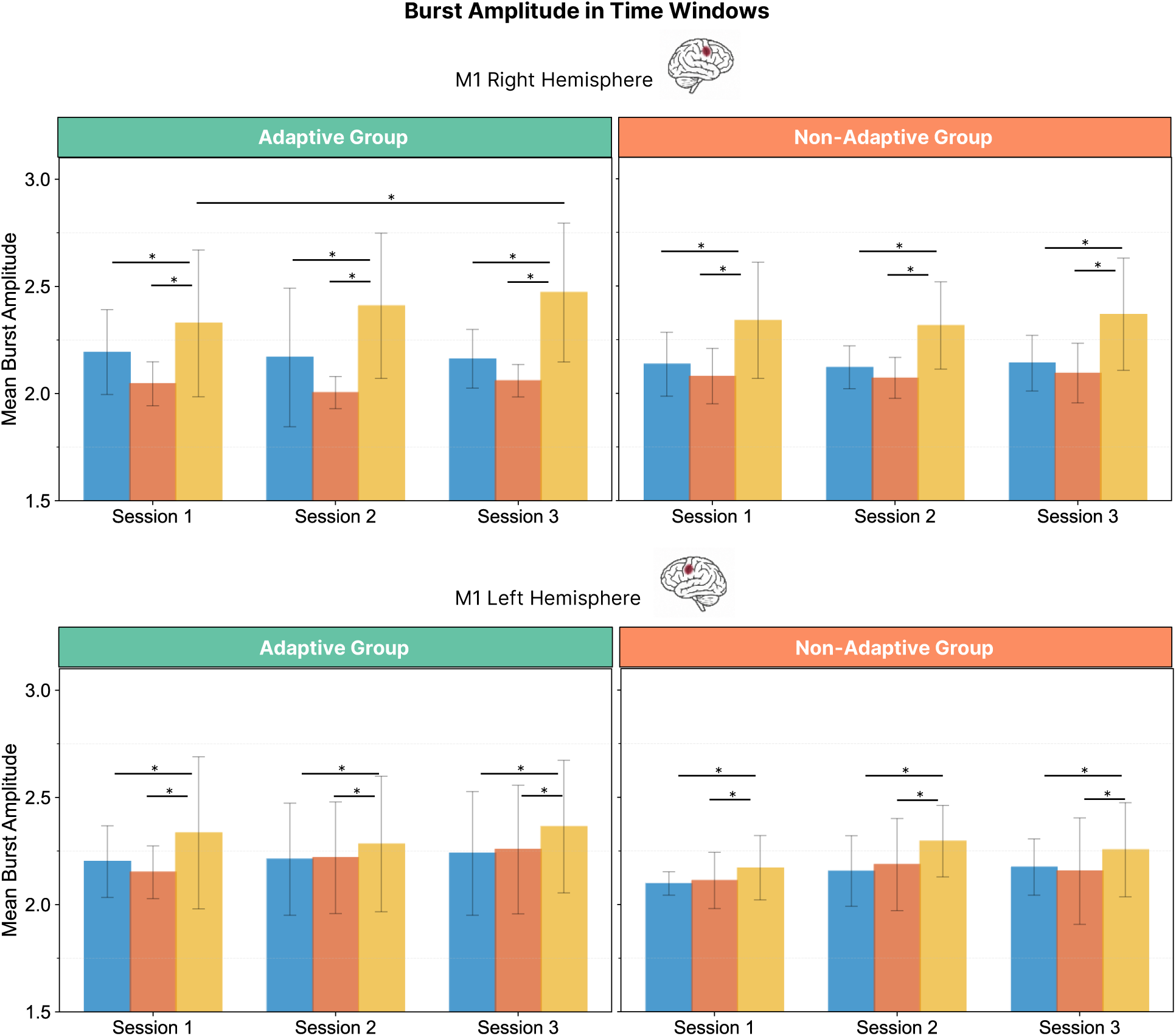
Beta burst amplitude across predefined time windows, and sessions. Average beta burst amplitude within each time window and session, plotted separately for right M1 (top panel) and left M1 (bottom panel). Asterisks indicate statistically significant differences between time windows (p < .05, Bonferroni-corrected).

**Figure 11.**
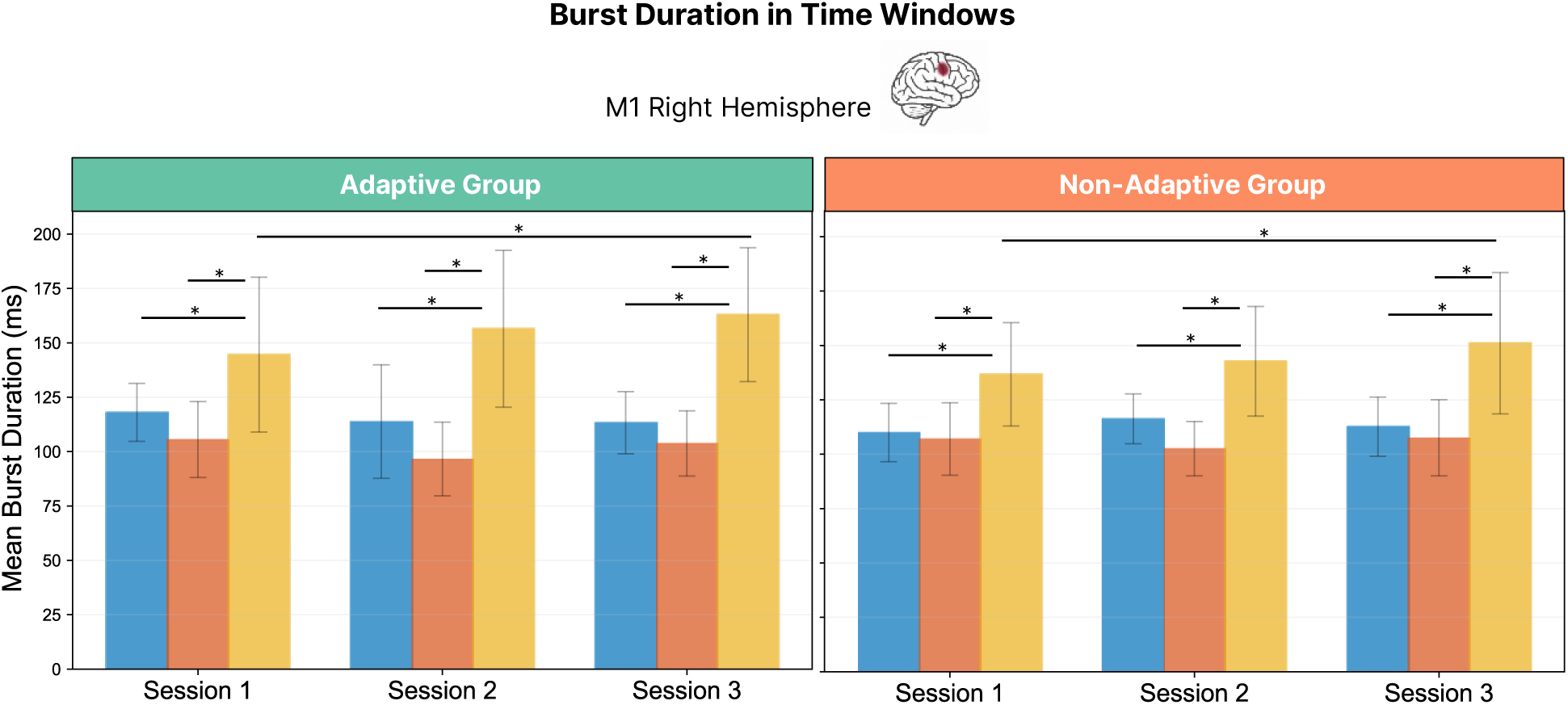
Beta burst duration across predefined time windows, and sessions. Average beta burst duration within each time window and session for right M1. Asterisks indicate statistically significant differences between time windows (p < .05, Bonferroni-corrected).

A repeated-measures ANOVA with Session as a within-subject factor and Group as a between-subject factor revealed a significant main effect of Session, F(2, 50) = 6.072, p = .004, indicating reduced trial-to-trial variability with training. Post hoc Bonferroni-corrected comparisons confirmed a significant reduction in variability between Session 1 and Session 3 (p = .011). The difference between Sessions 1 and 2 did not reach significance (p = .090), and no change was found between Sessions 2 and 3 (p = .659). No significant Session × Group interaction was observed, F(2, 50) = 0.459, p = .635, and the main effect of Group was also non-significant, F(1, 25) = 1.27, p = .270.

Next, we examined mean peak latency within the post-movement window. A repeated-measures ANOVA revealed a significant main effect of Session, *F*(1.38,34.44) = 7.11, *p* = .006 (Greenhouse–Geisser corrected). Across participants, mean peak latency shifted earlier with training, decreasing from Session 1 to both Session 2 (*p* = .025) and Session 3 (*p* = .030), with no difference between Sessions 2 and 3 (*p* = 1.000). Importantly, a significant Session × Group interaction was also observed, *F*(1.38,34.44) = 6.13, *p* = .011. Post hoc comparisons revealed that this effect was driven by the Non-Adaptive group, who showed later peaks in Session 1 and a marked shift to earlier latencies by Sessions 2 and 3 (Session 1 vs Session 2, *p* = .003; Session 1 vs Session 3, *p* = .044). In contrast, the Adaptive group showed more stable peak timing across sessions, with only a modest reduction that did not survive correction. By Session 3, both groups converged, with no significant difference in peak latency.

#### Training Increases Post-Movement beta burst amplitude in adaptive learners

We next examined whether beta burst amplitude also changed with training, focusing on peak burst amplitude within the three predefined time windows. A generalized linear mixed-effects model was applied to each M1 region separately, with fixed effects for Time Window, Session, and Group, and Participant included as a random intercept.

In right M1, burst amplitude varied significantly across task phases, as indicated by a main effect of Time Window, χ^2^(2) = 616.64, *p* < .001. Post hoc tests showed that amplitude was higher during Movement compared to Baseline (*p* = .021), and higher during the Post-Movement window compared to both Baseline (*p* < .001, Bonferroni-corrected) and Movement (*p* < .001). A main effect of Session was also observed, χ^2^(2) = 9.66, *p* = .008, reflecting changes over the course of training. Post hoc tests showed that amplitude increased significantly from Session 2 to Session 3 (*p* = .009, Bonferroni-corrected), while no significant differences were found between Sessions 1 and 2 (*p* = 1.000) or Sessions 1 and 3 (*p* = .056). Crucially, a significant Session × Time Window interaction emerged, (χ^2^(4) = 11.93, *p* = .018). Post hoc comparisons showed that in every session, Post-Movement amplitudes were higher than both Baseline and Movement (all *ps* < .001, Bonferroni-corrected). Moreover, within the Post-Movement window, amplitude increased significantly from Session 1 to Session 3 (*p* < .001), while no changes were detected in Baseline or Movement windows across sessions. A significant three-way interaction between Session, Time Window, and Group, was also found (χ^2^(4) = 13.27, *p* = .010). Within the Post-Movement, only the Adaptive group showed a significant increase across sessions: from 2.21 ± 0.49 in Session 1 to 2.23 ± 0.52 in Session 2 and 2.29 ± 0.53 in Session 3. This change reached significance between Session 1 and Session 3 (*p* < .001), but not between Sessions 1 and 2 (*p* = .412) or Sessions 2 and 3 (*p* = 1.000), suggesting a gradual training-related increase. In contrast, the Non-Adaptive group showed no significant change across sessions (2.18 ± 0.46, 2.19 ± 0.44, 2.22 ± 0.50; all *p*s =1.000). No significant training-related changes were found in the Baseline or Movement windows for either group, and there were no significant between-group differences at any single session.

In left M1, burst amplitude significantly differed across task phases, as indicated by a main effect of Time Window (χ^2^(2) = 633.94, p < .001). Post hoc tests showed that amplitudes were significantly higher during the Post-Movement window compared to both Baseline (p < .001, Bonferroni-corrected) and Movement (p < .001), while the difference between Baseline and Movement did not reach significance (p = .120). A significant main effect of Session was also observed (χ^2^(2) = 19.67, p < .001) reflecting changes across training. Post hoc tests revealed that amplitude increased significantly from Session 1 to Session 3 (p < .001, Bonferroni-corrected) and from Session 2 to Session 3 (p = .010), while no significant change was found between Sessions 1 and 2 (p = .497). Importantly, a significant Session × Group interaction was also detected (χ^2^(2) = 10.62, p = .005), reflecting divergent amplitude trajectories between the Adaptive and Non-Adaptive groups over time. In the Adaptive group, amplitude increased across training, with a significant rise from Session 1 to Session 3 (2.34 ± 0.64 vs. 2.43 ± 0.63; p = .001) and from Session 2 to Session 3 (2.36 ± 0.63 vs. 2.43 ± 0.63; p < .001), while no change was found between Sessions 1 and 2 (p = 1.000). In contrast, the Non-Adaptive group showed stable amplitudes across sessions (2.25 ± 0.52, 2.31 ± 0.50, 2.30 ± 0.54; all ps ≥ .236, Bonferroni-corrected). No between-group differences were detected at any single session (all ps = 1.000). Finally, the Session × Time Window interaction was not significant (p = .126), indicating that training-related changes were not specific to a particular task phase.

Burst amplitude and duration are inherently related: longer bursts provide more oscillatory cycles, which can increase the height of the envelope peak. Since we observed a significant Session × Time Window × Group interaction for burst amplitude in right M1, it was important to determine whether this effect could simply be explained by longer bursts. To address this, we ran a separate GLMM on burst duration in right M1. The analysis revealed a strong effect of task phase, χ^2^(2) = 476.52, p < .001. Post hoc tests confirmed that durations were longer during the Baseline window compared to Movement (p = .003, Bonferroni-corrected), and longest during the Post-Movement window, which exceeded both Baseline and Movement (both ps < .001). During Baseline and Movement, burst durations were on average ~100 ms, compared to 134–150 ms during Post-Movement. A significant Time Window × Session interaction was also observed, χ^2^(4) = 16.03, p = .003. Follow-up comparisons indicated that Post-Movement durations lengthened with training, increasing from 134 ms in Session 1 to 150 ms in Session 3 (p < .001), while Baseline and Movement durations remained relatively stable across sessions (~95–106 ms). No significant main effects of Session (p = .124) or Group (p = .351) were detected, and no higher-order interactions involving Group reached significance (all ps > .12). These results indicate that burst duration was consistently longer in the Post-Movement phase and showed a training-related increase in this window, but without group-specific effects.

## Discussion

We used a complex bimanual coordination task, practiced across multiple sessions, to track how beta-band activity changes during motor learning and whether these changes differ between two training approaches. We expected participants to show general improvements in bimanual coordination across sessions, with greater and more sustained gains in the Adaptive group due to the incremental demands of difficulty scaling. Beta changes were strongest in right sensorimotor cortex, which guided our focus on bilateral M1 for the burst-resolved analyses. Based on this, we tested three main neural hypotheses: (1) ERD/ERS patterns would become more pronounced with practice; (2) these average power changes would reflect clearer temporal segregation of beta bursts across task phases; and (3) adaptive training would lead to stronger changes in burst dynamics due to its sustained performance demands.

### Optimizing Motor Learning Requires Well-Timed, Individualized Challenge

Both training protocols led to significant improvements in bimanual coordination from baseline to retention, demonstrating effective learning across groups. However, participants in the adaptive condition showed significantly greater gains during training. This aligns with the Challenge Point Framework (Guadagnoli & Lee, 2004), which suggests learning is maximized when task demands are matched to the learner’s ability. Properly calibrated challenge makes errors more informative, boosts learners’ ability to adjust, and sustains motivation. The adaptive protocol likely maintained this optimal learning zone by continuously adjusting difficulty, promoting deeper engagement and faster improvement.

Despite these stronger gains during training, the adaptive group showed a small but significant decline in retention performance. This seems to contradict the principle of desirable difficulty (Bjork & Bjork, 2011), which argues that challenging practice supports long-term retention. A likely explanation is that in our adaptive protocol, task difficulty increased immediately once participants reached 80% accuracy, leaving little time to stabilize skills within each new tolerance range before the difficulty was raised again. This rapid progression may have accelerated adaptation but limited opportunities for consolidation, weakening long-term retention.

Conversely, the non-adaptive condition, though designed as a control, may have provided a more favourable overall difficulty. With task constraints fixed, some participants faced a sustained mismatch between their abilities and task demands. This persistent challenge aligns with Lövdén’s mismatch-driven plasticity framework and the principle of desirable difficulty (Bjork & Bjork, 2011) where extended struggle promotes deeper consolidation. Ironically, while the adaptive group progressed more quickly during training, some members of the non-adaptive group may have benefited from a more challenging yet ultimately more stabilizing learning environment.

### Training Modulates Beta Burst Probability and Sharpens Temporal Precision

With training, beta activity showed stronger desynchronization during movement (ERD) and increased synchronization after movement (ERS), primarily in the right sensorimotor cortex. Source-level analysis localized these changes to the right primary motor cortex (M1), contralateral to the active hand. This aligns with a large body of animal work, showing that M1 not only generates motor output but also dynamically adapts its local activity with learning, integrating subcortical and cortical inputs in a use-dependent manner (Kawai et al., 2015; Peters et al., 2017). Such modulation provides compelling evidence for learning-related reorganisation within human motor cortex, supporting the consolidation of connections associated with the acquired skill (see also (Pascual-Leone et al., 1995).

At a physiological level, these changes are likely driven by shifting thalamocortical input patterns: at rest, the thalamus provides tonic excitatory drive to M1, supporting intermittent beta bursts that reflect a stable, inhibited motor state. During movement preparation and execution, this input becomes more phasic and less synchronized, reducing burst occurrence and allowing the cortex to enter a more excitable, movement-ready state (Shepherd & Yamawaki, 2021). Learning may refine this balance, leading to more efficient modulation of beta bursts in M1 to support flexible motor control and movement termination (Ramot et al., 2025). These observations are also consistent with rodent studies showing that acquisition of a grasping skill selectively strengthens intracortical connections (i.e. increased cortical excitability) within the “trained” M1 regions, while the “untrained” homologous regions remained unchanged (Rioult-Pedotti et al., 1998). Further investigations are warranted to elucidate whether modulations of beta bursts’ dynamic are mechanistically associated with alterations in horizontal connections within M1 regions.

To determine whether motor learning alters the structure of beta bursts themselves, we examined changes in their probability, timing, and amplitude across training. The most pronounced learning-related shifts were observed in burst probability and timing, reflecting increased selectivity and temporal precision in motor-related neural activity. In contralateral M1, during movement, bursts occurred less frequently relative to baseline, particularly in later sessions, suggesting a refinement of inhibitory control mechanisms in support of more phasic and precisely timed force output. By contrast, post-movement bursts became more frequent and more tightly time-locked to movement offset, indicating more consistent engagement of evaluative or inhibitory processes (Spitzer & Haegens, 2017; Tan et al., 2014). Interestingly, the Non-Adaptive group showed later post-movement burst peaks early in training, which could hypothetically reflect more variable force release dynamics—such as some individuals sustaining their motor output longer before fully terminating the action (see Fig. 3). Although we did not directly quantify this effect, it suggests that post-movement bursts may be tightly coupled to the precision of movement offset. More generally, the increasing reliability of these bursts suggests that post-movement bursts mark the successful completion of an action, acting as a neural “full stop” that terminates motor output and prepares the system for the next action (Lundqvist et al., 2024). Over time, these bursts grew more stereotyped and temporally sharpened, likely reflecting the consolidation of internal control signals and a shift from feedback-driven corrections to feedforward control based on stable internal models (Tan et al., 2016; Barone et al., 2021). These refinements appear to reflect a general feature of motor learning, consistent across training conditions. In line with the Exploration–Selection–Refinement (ESR) model (Wenger et al., 2017), they point to a progression from variable, exploratory responses to stable and precisely timed neural patterns that support skilled performance.

In this study, we focused on beta bursts localized to the hand area of bilateral M1, based on the spatial specificity of our source-level findings and our hypotheses. However, beta dynamics in M1 are likely shaped by a broader cortical network. Regions such as the supplementary motor area (SMA), premotor cortex, and prefrontal cortex—involved in movement planning, sequencing, and performance monitoring—may influence M1 through top-down projections that modulate the timing and structure of local beta bursts (Adams et al., 2013; Khanna & Carmena, 2015; Narayanan & Laubach, 2006; Weiler et al., 2008). To fully understand how motor learning reorganizes these dynamics, it is essential to examine how bursts unfold across a wider network. For example, determining whether bursts in SMA or premotor cortex systematically lead, lag, or synchronize with those in M1 could reveal how cortical areas coordinate motor preparation and evaluation. In bimanual task, like this one, interhemispheric interactions are also likely involved. While only right M1 showed a significant increase in post-movement burst probability across sessions, a similar, though non-significant, trend was observed in left M1. Raster plots suggest that phasic left-hand movements modulate beta activity in both hemispheres, potentially reflecting bilateral coordination or cortical crosstalk (Cardoso de Oliveira et al., 2001; Daffertshofer et al., 2005; Gerloff & Andres, 2002; Liuzzi et al., 2011; Rueda-Delgado et al., 2017). Because beta activity emerges as brief, transient bursts, rather than sustained oscillations, traditional phase-based connectivity metrics may miss these fine-grained dynamics. Analyses based on burst timing and co-occurrence may offer a more mechanistic, event-based perspective on how distributed motor circuits interact and adapt through learning (Seedat et al., 2020b).

### Adaptive Training Amplifies Beta Burst Strength?

While ERD and ERS primarily reflect changes in beta burst probability, we also examined whether burst amplitude changed with training regimen. Amplitude varied systematically across task phases, being lowest during movement, intermediate at baseline, and highest post-movement. This pattern likely reflects state-dependent dynamics within M1: during movement, local beta synchrony is suppressed to facilitate motor output, limiting burst strength; at baseline, activity is relatively stable but not strongly engaged; and after movement, rebound-related processes such as inhibitory re-engagement and synchronized thalamocortical input create conditions for higher-amplitude bursts. Only the adaptive group, however, showed a significant increase in post-movement burst amplitude in contralateral M1, indicating a training-related change in the underlying activity patterns generating these bursts.

Amplitude reflects the strength of synchronous neural activity, but it is also shaped by multiple morphological properties of the burst itself, including its duration, waveform shape, and spectral composition (Cole & Voytek, 2017; Sherman et al., 2016). For example, longer bursts can elevate the envelope peak simply by summing more cycles, while bursts narrowly confined to the beta range may reach a higher peak than those with broader frequency spread. Thus, amplitude should not be treated as a straightforward proxy of cortical drive, but rather as one complementary dimension of burst structure, alongside probability and timing.

Given that one of the main morphological factors influencing amplitude is burst duration, we next tested whether the observed amplitude effect could simply reflect longer bursts. If this were the case, the Adaptive-specific increase in post-movement amplitude would be explained by corresponding group differences in duration. However, burst duration—although consistently longest in the post-movement phase and lengthening modestly with practice—showed similar changes across groups and therefore could not account for the Adaptive-specific amplitude increase.

Within this broader context, the Adaptive-specific increase in post-movement amplitude can be interpreted as reflecting stronger and more coherent synaptic input to pyramidal neurons, particularly through the coordinated timing of proximal excitation and distal inhibition (Bonaiuto et al., 2021; Sherman et al., 2016; West et al., 2023). This synchrony increases transmembrane current flow, producing larger beta-band deflections in the source signal. More coherent input may reflect a more stable local state in M1 after movement completion.

At the network level, increased burst amplitude may reflect a reduced influence of top-down or corrective input. In early learning, feedback-related activity from prefrontal, premotor, or subcortical regions may disrupt synchrony in M1. With practice, as internal models become more accurate, M1 may operate with greater autonomy, receiving more consistent thalamocortical input and producing stronger, more uniform bursts. This amplitude increase was specific to the adaptive group, who were required to continually adjust their performance across sessions. Repeated adaptation likely drove ongoing updates to internal models. By the end of training, these participants may have developed more accurate and stable predictions, allowing for cleaner and more synchronized inputs to M1, reflected in increased burst amplitude. These findings suggest that burst amplitude may serve as an index of internal model refinement during adaptive motor learning.

### Methodological considerations

EEG provides limited access to subcortical activity, even though regions such as the thalamus, basal ganglia, and cerebellum play key roles in motor learning. We therefore centered our analyses on M1 for two reasons: first, both scalp-and source-level findings consistently highlighted the strongest beta changes in right sensorimotor cortex; and second, M1 is a well-characterized and accessible target that is tightly embedded in subcortical loops, making it a robust cortical anchor for studying learning-related beta bursts.

## Conclusion

This study highlights that primary motor cortex (M1) is not merely an output structure issuing motor commands, but a dynamic site of input integration and experience-dependent reorganization. Across training, our data-driven approach reveals that learning-related changes in beta activity were most evident in contralateral sensorimotor regions, particularly in right M1, suggesting that this area may play a key role in reorganizing motor control with practice. Using source-resolved EEG and burst-level analysis, we found that motor learning shapes beta activity in both timing and strength. With practice, beta bursts in M1 became more precisely time-locked to task phases, particularly after movement, reflecting more reliable and task phase-specific modulation. These timing refinements were observed across both training protocols, suggesting they represent a general feature of motor learning. Under adaptive training, beta bursts also increased in amplitude. This may reflect more synchronized synaptic input to pyramidal neurons in M1, potentially due to reduced interference from variable top-down signals. One interpretation is that increased burst amplitude relates to the refinement of internal models, as learners rely more on predictive control and less on feedback-driven correction. However, this interpretation remains tentative and should be explored in future studies.

## Materials and Methods

### Participants

Thirty-two right-handed university students (16 females; mean age = 21 ± 2.46 years) participated in the study. All had normal or corrected-to-normal vision, no neurological or psychiatric history, and gave written informed consent. The protocol was approved by the Vaud Cantonal Ethics Committee (2022-01653). One participant was excluded due to task misunderstanding, and four others were removed due to EEG artifacts caused by muscle contractions or movement at the moment of the left-hand shot, yielding a final EEG sample of 27.

### Experimental procedure

Over eight weeks, participants completed 13 sessions: two MRI scans (Sessions 1 and 12), one baseline (Session 2), nine training sessions (Sessions 3–11), and one retention test (Session 13). EEG was recorded during Sessions 3, 7, and 11 (i.e., training Sessions 1, 5, and 9). Participants were randomly assigned to either an adaptive or non-adaptive training group (see Fig. 1-A). Each training session consisted of 60 trials (12 blocks of 5 trials), with breaks between every 4 blocks. Baseline and retention sessions consisted of 30 consecutive no-feedback trials (see Fig. 1-B).

### Bimanual Motor Task

Participants performed a visually guided bimanual force coordination task while seated in a dimly lit room, with both arms resting on a custom support (Fig. 2-A). They held dynamometers in each hand (BIOPAC, TSD121C), gripping at standardized positions. Task force was set at 25% of each participant’s lower maximum voluntary contraction (MVC).

Each trial began with the right hand maintaining a steady force at 25% MVC (±2%) for 300 ms, guided by real-time visual feedback indicating whether the force was within range. Once stabilized, a static on-screen cursor began a 3-second circular trajectory at constant speed. Cursor movement was time-locked and independent of force. Participants were instructed to maintain the right-hand force throughout the trial without further feedback. A single visual target was positioned at a random point along the path (between π/4 and 2π − π/4 radians). When the cursor reached the target, participants applied a brief, ballistic left-hand force—also at 25% MVC—to “shoot” it. This created a bimanual challenge requiring both sustained force control (right hand) and precisely timed ballistic responses (left hand).

After each trial, participants received visual feedback indicating whether their performance for each hand fell within the predefined tolerance criteria (Fig. 2C). Right-hand performance was considered correct if force remained within the target range for the entire 3-second trajectory. Left-hand performance was correct if the ballistic force fell within range and occurred within ±125 ms of the target center. Augmented summary feedback was provided every five trials in the first four sessions and every ten trials in the last five sessions. This feedback displayed two key metrics: (1) the percentage of the 3-second trajectory during which right-hand force was valid, and (2) the percentage of successfully destroyed targets. These intervals were based on prior motor learning research demonstrating optimal feedback frequency for skill acquisition (G et al., 1993; Lai & Shea, 1998; Winstein & Schmidt, 1990).

The retention session followed the same procedure but was performed without feedback to assess learning.

### Training Protocol

Participants were assigned to one of two training groups—adaptive or non-adaptive—after completing the baseline motor task. Both groups performed the same bimanual force task, but with different difficulty adjustments. In the adaptive group, task difficulty increased based on individual performance across sessions. Specifically, if accuracy for either hand was ≥80%, its tolerance range was narrowed for the next session to include only the 70% of responses closest to the target force from the previous session. If accuracy was <80%, the tolerance range remained unchanged. In the non-adaptive group, difficulty remained fixed using pilot-derived tolerance values, set such that ~50% of baseline responses fell within range.

Tolerance ranges defined the allowable margin of error around the target force, separately for each hand (cursor and target). Narrower ranges increased difficulty by demanding greater force precision. To ensure balanced groups, participants were matched by age, sex, and baseline performance using RMSE comparisons. All sessions were conducted at the Experimental Behavioural Research Laboratory (LERB), University of Lausanne.

### EEG Data Acquisition and Preprocessing

EEG data were recorded during training sessions 1, 5, and 9 (protocol Sessions 3, 7, and 11) using a 128-channel Biosemi ActiveTwo system (Biosemi, Amsterdam, The Netherlands), at a sampling rate of 1024 Hz. Electrode positions were digitized using the NDI Krios scanner. Signals were downsampled to 256 Hz, high-pass filtered at 1 Hz, and notch-filtered at 50 Hz. Independent Component Analysis (ICA) was used to remove artifacts corresponding to eye movements, muscle activity, or other physiological noise via the Multiple Artifact Rejection Algorithm (MARA).

Components explaining more than 90% of the variance were removed (mean number of components removed: 23 ±9). Bad channels were interpolated, and data were re-referenced to the average. Epochs were extracted from −2000 to +2000 ms around the left-hand force peak. Five participants were excluded due to excessive artifacts, yielding a final EEG sample of 27.

### MRI Data Acquisition and Preprocessing

MRI data were collected on a 3 T Siemens Magnetom Prisma scanner (Siemens Healthcare, Erlangen, Germany) equipped with a 64-channel head–neck coil, and included T1-weighted (T1w), T2-weighted (T2w), multi-parameter mapping (MPM)(1.5 mm isotropic) and diffusion-weighted imaging (DWI)(2.2mm isotropic) sequences.

For preprocessing we used a semi-automated MATLAB pipeline based on Brainstorm. Fiducial points (nasion, left/right pre-auricular) were manually labeled on the T1w image. Anatomical segmentation via CAT12 generated masks of scalp, skull, and brain compartments, from which individualized Boundary Element Model (BEM) surfaces were derived for EEG source modeling.

### EEG-MRI Coregistration

EEG sensor locations were coregistered to each participant’s MRI by aligning digitized fiducials (nasion, left/right pre-auricular points) with corresponding MRI landmarks. This alignment was refined using an iterative rigid-body transformation minimizing the distance between digitized scalp points and the MRI-derived scalp surface. Final sensor positions were projected onto the scalp mesh for accurate forward model construction.

### EEG data analysis

#### Time-Frequency Analysis

To examine training-related changes in beta activity, we computed time–frequency representations (TFRs) from −0.75 to 1.5 s around movement onset to remove any edge effects. TFRs were generated per subject and session using wavelet convolution in FieldTrip (Oostenveld et al., 2011), covering 15–30 Hz in 1 Hz steps. A custom MATLAB function (Delorme & Makeig, 2004) implemented a nonlinear scaling of wavelet cycles to optimize the trade-off between temporal and frequency resolution. For each time–frequency point, power was normalized to the session-specific mean (relative baseline correction), enabling comparability across sessions.

Session differences (Sessions 1 vs. 3) were evaluated using non-parametric cluster-based permutation tests (Maris & Oostenveld, 2007) implemented in FieldTrip. The analysis was based on 1000 random permutations of a dependent-samples T-statistic. Clusters were formed using a threshold of p < 0.025 and a minimum of two neighbouring electrodes (defined as ≤30 mm apart, based on the Biosemi128 layout). Cluster-level significance was assessed via two-tailed Monte Carlo correction (α = 0.05). For follow-up visualization and statistics, the three electrodes with the highest average T-values within the significant cluster (B21, B22, B23) were extracted.

#### Source-level Analysis

Cortical sources were reconstructed using weighted minimum norm estimation (wMNE) in Brainstorm (Tadel et al., 2011). For each participant, we used individualized BEM head models derived from their MRI segmentations and digitized electrode positions (see Fig.12). The inverse solution was computed on a cortical mesh of 15,002 vertices assuming fixed orientations, with depth weighting (exponent = 0.5; weight limit = 10), a signal-to-noise ratio (SNR) of 3, and a fixed noise level of 0.1. Subject-and session-specific source kernels were then applied to preprocessed epochs to extract trial-wise source-level time series.

**Figure 12.**
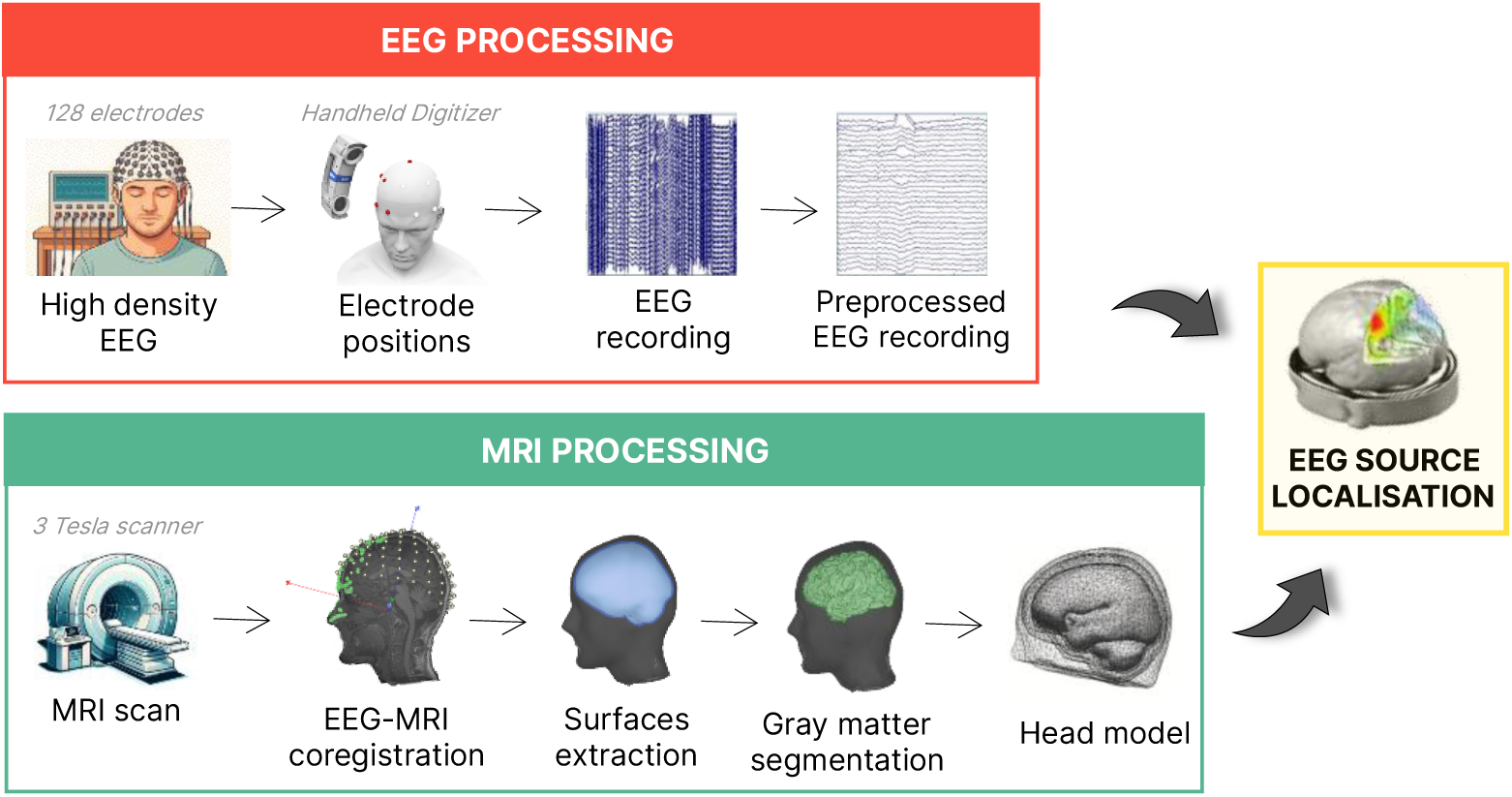
EEG and MRI processing pipeline for source localization. EEG data were recorded using 128 electrodes and spatially coregistered with individual head anatomy using a handheld digitizer. MRI data were acquired using a 3 Tesla scanner to reconstruct cortical surfaces. EEG and MRI data were combined to perform source localization and estimate cortical sources of EEG activity.

Beta-band power was estimated by bandpass filtering the source time series (15-30Hz), applying the Hilbert transform to compute instantaneous power, and averaging across trials. Power maps were normalized per vertex by dividing by the mean activity from −1 to 1 s. These maps were projected onto the ICBM152 cortical surface to allow group-level comparisons.

Session-related changes in beta power were evaluated using paired cluster-based permutation tests separately for each group. Cluster paired t-tests compared Sessions 1 and 3 within two a priori time windows: movement (−400 to +250 ms) and post-movement (250 to 1000 ms). Spatial clusters were defined by vertex adjacency, with a cluster-forming threshold of p < 0.005. Cluster-level significance was assessed using Monte Carlo correction (α = 0.05) based on 1000 permutations.

#### Beta Burst Detection

To investigate session-related changes in transient beta activity, we focused on the primary motor cortex (M1), guided by source-level contrasts spanning the broader sensorimotor regions (S1 and M1). While beta bursts are not uniquely motor-related, our hypotheses targeted their role in motor control. We therefore restricted burst detection to M1, the principal cortical generator of movement-related beta dynamics to test whether motor training reorganizes bursts in a functionally specific manner.

For each participant, we defined individualized left and right M1 hand areas scouts (~140 vertices centered on MNI [38, −22, 54]) to ensure anatomically consistent sampling. Source time series were bandpass-filtered around each subject’s peak beta frequency (±3 Hz), and the amplitude envelope was extracted via Hilbert transform. Envelopes were normalized per trial relative to session means.

Burst detection thresholds were optimized using a data-driven procedure adapted from Little et al., (2019). Multiple thresholds—defined as SDs above the median amplitude—were tested. For each, we calculated the number of bursts per trial and computed its Spearman correlation with mean trial amplitude. The SD multiple yielding the highest average correlation across all subjects, sessions, and scouts was selected and applied consistently across the dataset across all subjects and sessions (see Fig.13).

**Figure 13.**
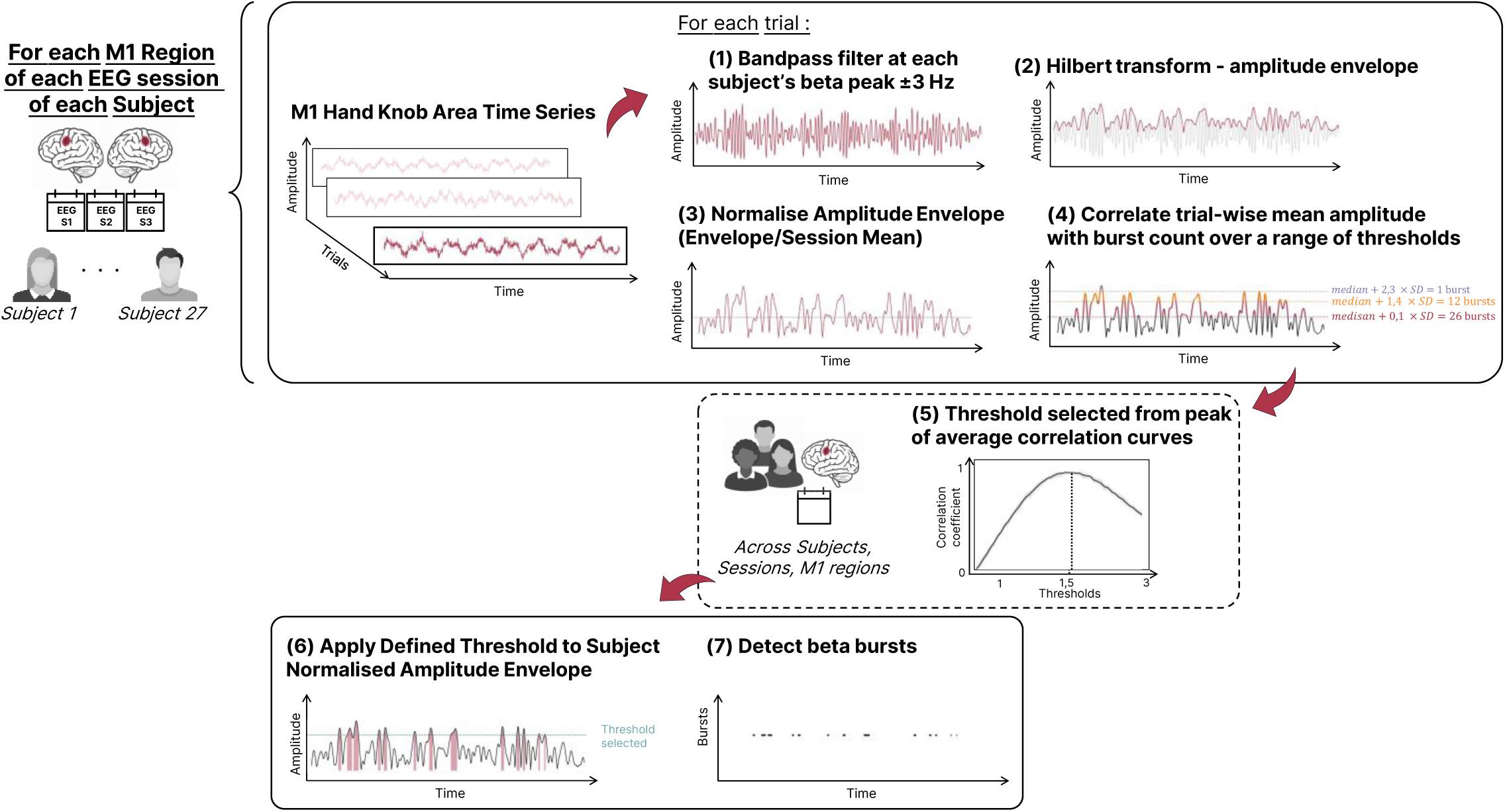
Beta burst detection pipeline. Source-level time series from individualized M1 scouts were filtered around each subject’s peak beta frequency (±3 Hz), and the amplitude envelope was extracted using the Hilbert transform and normalized per trial. Detection thresholds were tested as multiples of the standard deviation above the median amplitude. For each threshold, the number of suprathreshold bursts per trial was computed and correlated with mean trial amplitude using Spearman correlation. Correlation coefficients were averaged across subjects, sessions, and M1 regions (dotted panel), and the threshold yielding the highest average correlation was selected and then applied uniformly to each subject’s data to detect beta bursts.

#### Beta Burst Metrics

We examined three complementary burst metrics across three trial-aligned windows— Baseline (−1750 to −750 ms), Movement (−750 to 250 ms), and Post-movement (250 to 1250 ms)— all time-locked to the left-hand shot.

Burst probability was defined as the proportion of suprathreshold time points per trial, determined by binarizing the beta amplitude envelope at each time point (1 = above threshold, 0 = below). These binary matrices were averaged across trials and scaled by the total number of trials, yielding a time-resolved measure of burst likelihood per subject, session, and scout. This metric reflects how consistently bursts occur during distinct task phases.

Burst timing variability was used to assess the consistency of post-movement beta burst timing across trials. For each trial, the binary burst matrix was smoothed using a Gaussian kernel, and the first peak in the smoothed signal occurring after time zero was detected. This yielded a distribution of post-zero burst peak times across trials. For each subject and session, we computed the standard deviation of these peak times as a measure of timing variability, where lower values indicate more temporally consistent burst occurrence following movement execution.

Burst amplitude was defined as the average of all burst peak amplitudes occurring within each predefined time window, across trials.

