## Supplementary Material for "Beta-burst dynamics in the motor cortex are reshaped through sensorimotor refinement"

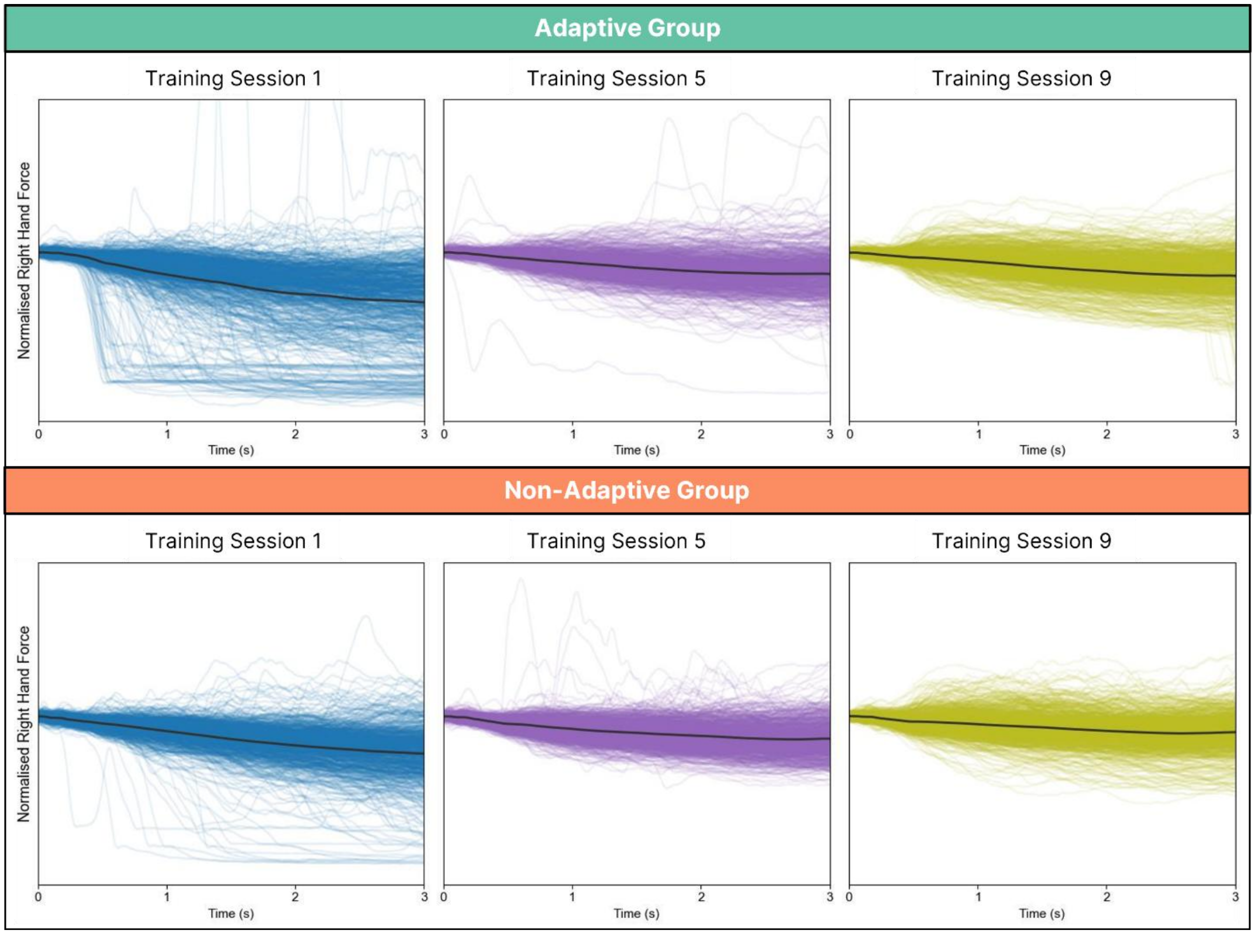


**Figure S1 | Normalized right-hand force profiles across training sessions.**
Force traces for Sessions 1 (blue), 5 (purple), and 9 (green) are shown for the adaptive group (top row) and non-adaptive group (bottom row). Thin lines represent individual trials from all participants; thick black lines indicate the average across trials. Force was normalized within each trial, and traces are plotted over the full 3-second movement period (0–3 s), during which participants were instructed to maintain steady right-hand force.

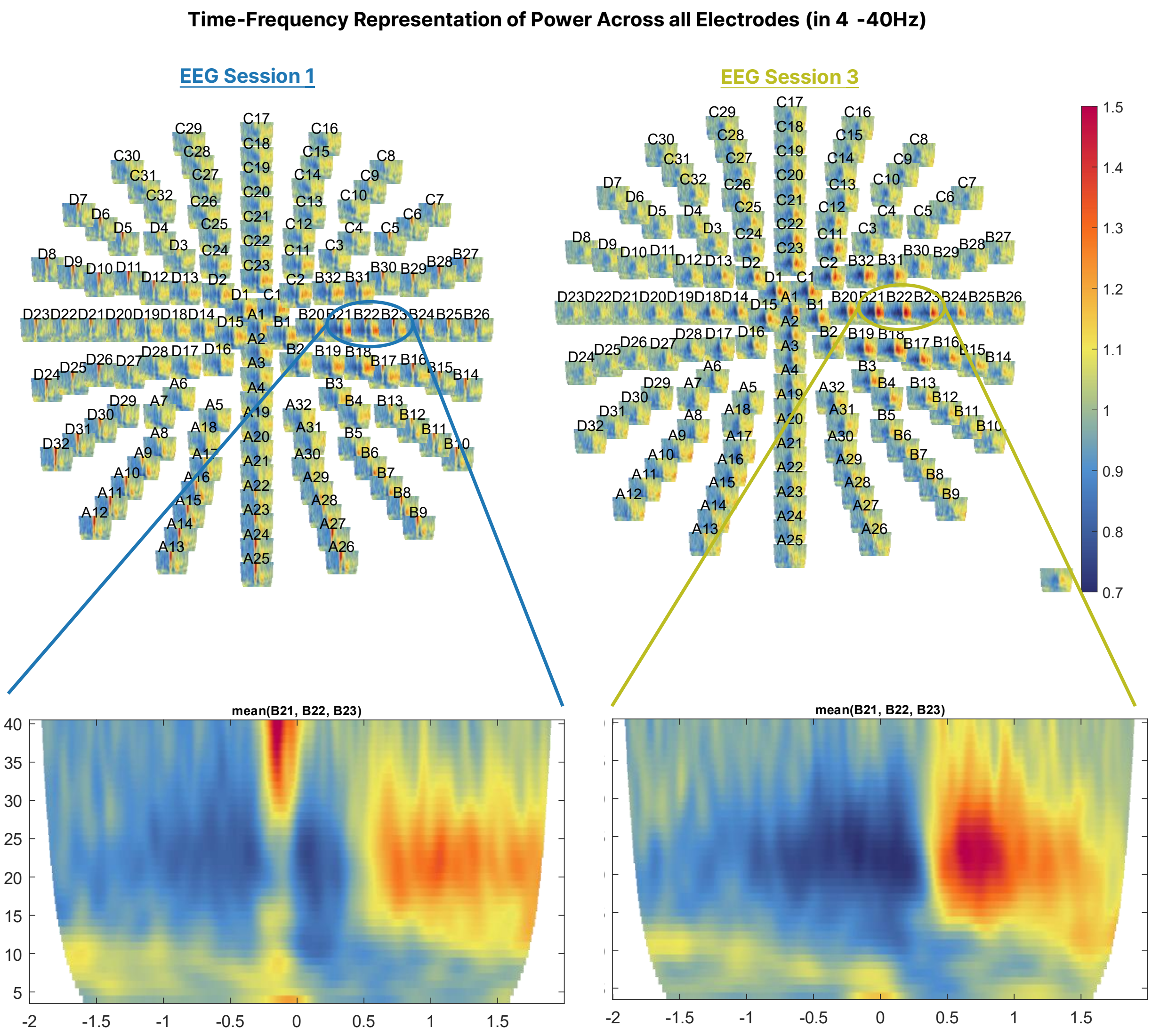


**Figure S2 | Session-wise changes in broadband-band activity at the sensor level.**
Scalp topographies show the spatial distribution of relative beta power (4–40 Hz) during the bimanual time period for EEG Sessions 1 (blue) and 3 (green). Each subplot represents the full sensor array, with warmer colors indicating higher power. Bottom panels display time–frequency representations averaged across three centro-parietal electrodes (B21, B22, B23), which were later identified as showing the strongest training-related effects in the beta band.

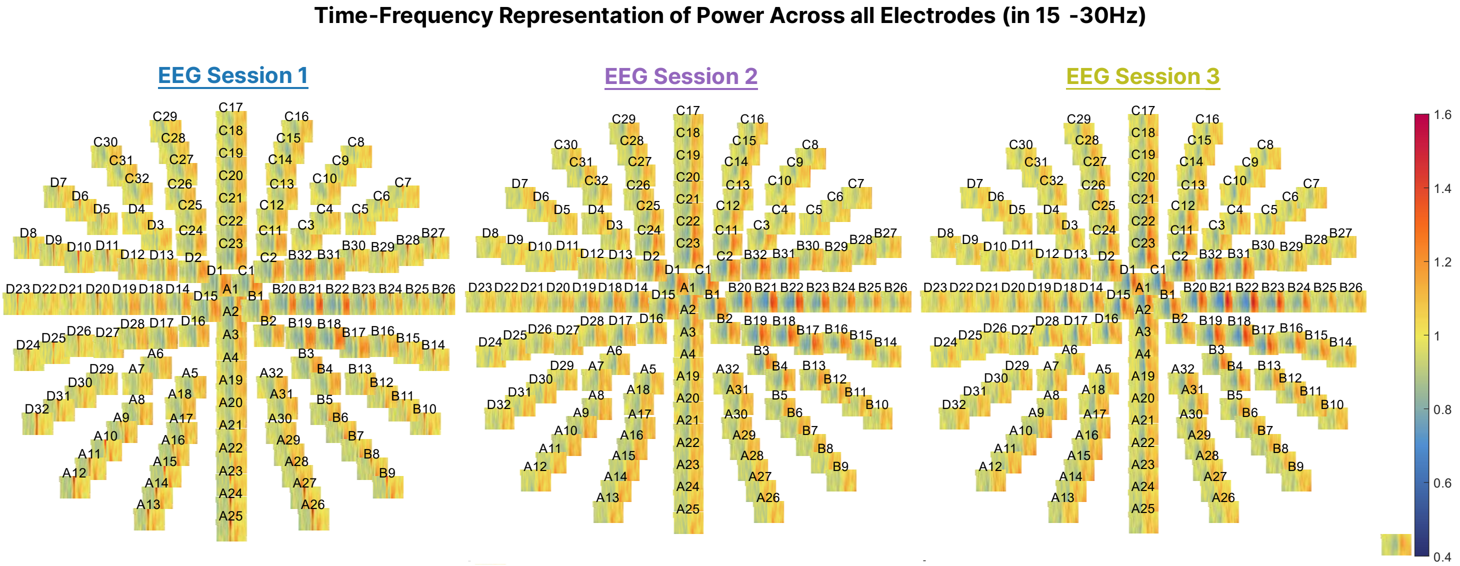


**Figure S3 | Session-wise changes in beta-band activity at the sensor level.**
Scalp topographies show the spatial distribution of relative beta power (15–30 Hz) during the bimanual time period for EEG Sessions 1 (blue), 2 (purple), and 3 (green). Each subplot represents the full sensor array, with warmer colors indicating higher power

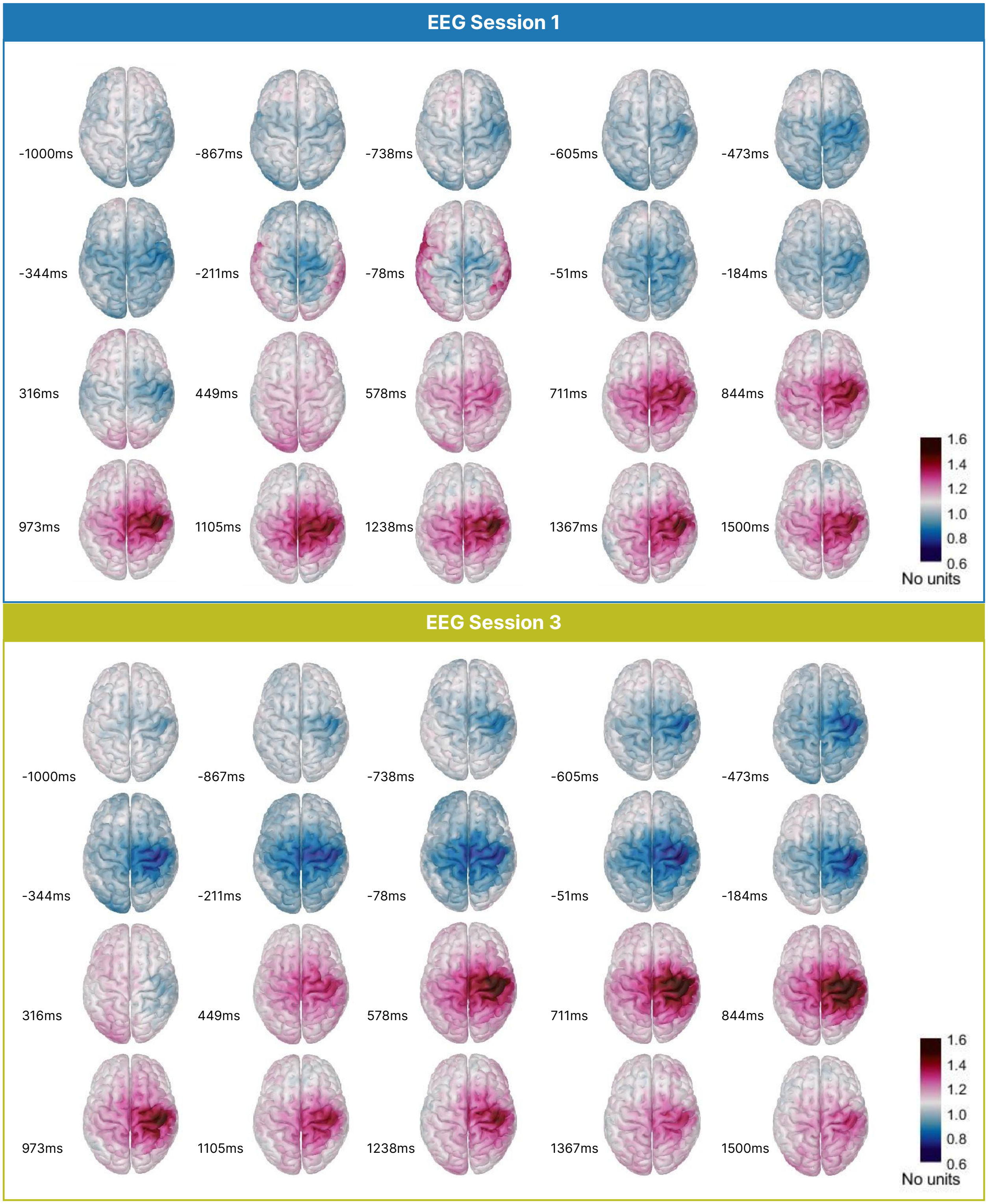


**Figure S4 | Source-level maps of relative beta-band power during the bimanual phase in Sessions 1 and 3.** Dorsal cortical maps of relative beta power (15–30 Hz) during the bimanual coordination window (–250 to 1250 ms relative to movement onset) for EEG Sessions 1 (top) and 3 (bottom). Power is expressed relative to the average across the full trial; values below 1 (blue) indicate desynchronization, and values above 1 (pink to dark red) indicate synchronization.

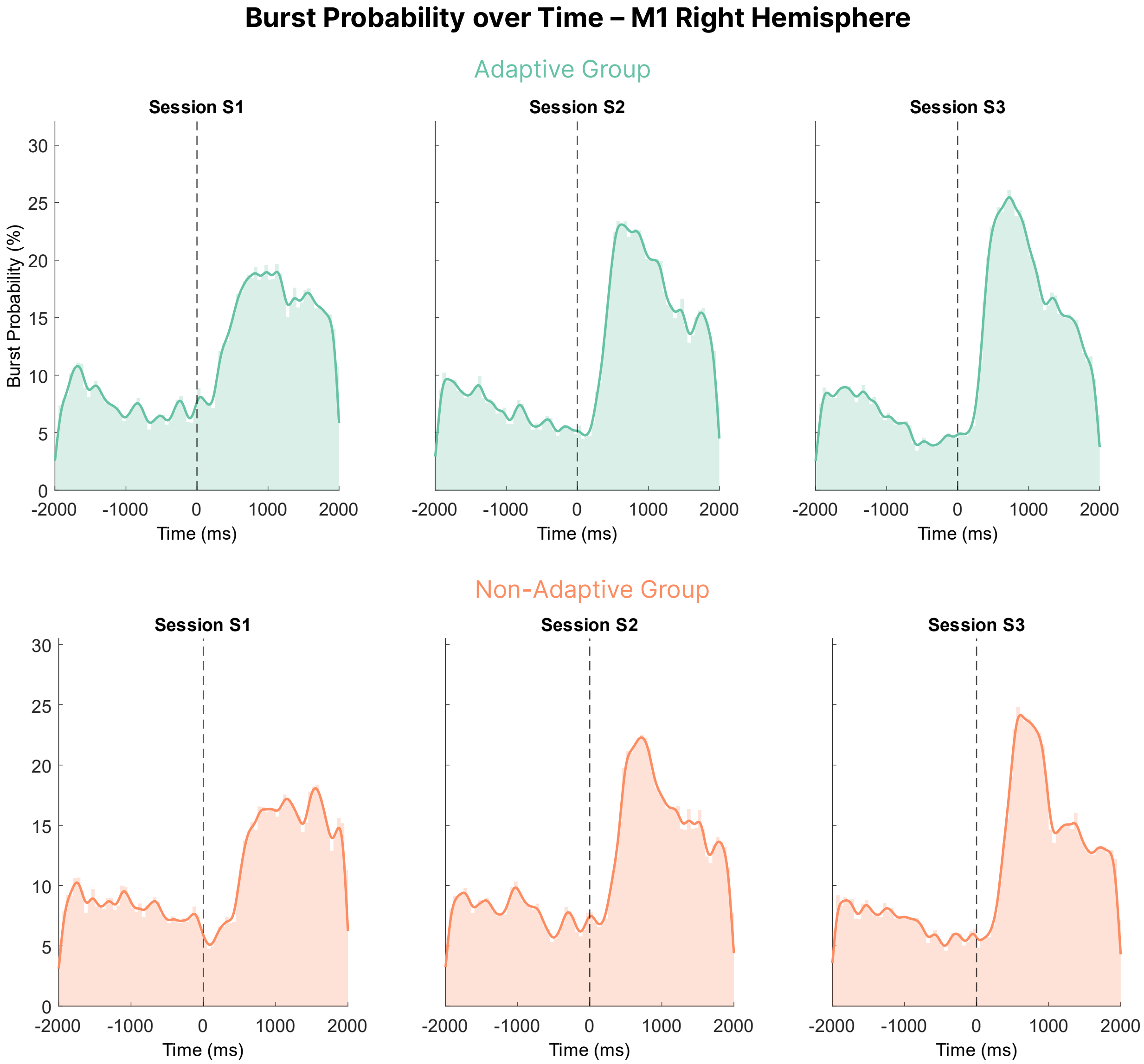


**Figure S5 | Time-resolved beta burst probability across training sessions in right M1.** Group-level beta burst probability (% of trials with a burst) is plotted over time relative to movement onset (dashed line at 0 ms), for the Adaptive group (left, teal) and Non-Adaptive group (right, orange). Each panel shows data from one EEG session (S1–S3). Shaded bars indicate average burst probability in 50 ms bins; solid lines reflect time courses smoothed with a Gaussian kernel ($\sigma = 50$ ms).
